# Ecological cues orchestrate concerted courtship in a *Drosophila* host specialist

**DOI:** 10.1101/2025.10.13.681913

**Authors:** Philipp Brand, Katherine Keller, Rory T. Coleman, Noelle B. Eghbali, Sarah Zylka, Lucia Prieto-Godino, Vanessa Ruta

## Abstract

Mating decisions are often attributed to the sensory signaling between prospective sexual partners. Yet these interactions are also shaped by the broader environmental context in which they unfold, to appropriately align sexual arousal with reproductive opportunities. Here we show that in the host specialist *Drosophila erecta* mating is strictly contingent on the ecological and social environment generated as flies densely aggregate in groups on a food patch. We find that food volatiles directly promote male sexual arousal, triggering individuals to sample and pursue potential mates, giving rise to dynamic interactions across the group. The ensuing visual motion transforms each male’s visual field, which in turn further amplifies his arousal, generating a multisensory feedback loop that coordinately promotes courtship across individuals. *D. erecta’s* strict dependence on environmental cues appears latent in related species, such as *D. melanogaster,* where food odor can promote arousal but is dispensable for vigorous courtship. Comparative circuit analyses reveal that species-specific thresholds for sexual arousal reflect variation in how olfactory input modulates conserved nodes controlling courtship drive, rendering food volatiles a strict sensory gate only in *D. erecta*. Together, our findings highlight how ecological cues not directly tied to sexual signaling can profoundly influence reproductive behavior and reorganize the social landscape to ensure mating occurs in contexts where reproductive opportunities are abundant.

---

Social cues are essential for coordinating interactions between individuals, facilitating the emergence of group dynamics including mating, aggression, and collective foraging^1–4^. Yet social interactions do not occur in isolation. Rather, they unfold within a broader environmental context that structures when, where, and how social behaviors are expressed. Courtship and mating, for example, often take place at ecologically privileged locations— food sources^5–7^, prominent visual landmarks^8–11^, or suitable nesting sites^12–15^—where many individuals aggregate, either to increase their encounters with prospective sexual partners or to access key resources that promote reproductive success. In these contexts, a central component of the environment is the social collective itself— the multisensory backdrop that emerges from the movement, sensory signals, and interactions across all individuals within a group^17–19^. Social behaviors are therefore likely to be shaped by both the static properties of the physical environment and the dynamic features of the social landscape. Yet the neural mechanisms that integrate these distinct sensory streams to regulate social arousal states and structure interindividual interactions remain largely unknown.

The mating behaviors of *Drosophila* offer a powerful inroad to explore how animals adapt to complex ecological and social landscapes. In the wild, flies meet and mate on their host plants, typically decaying ephemeral substrates where many individuals congregate^7^. Members of the *Drosophila* genus vary widely in their ecology, ranging from generalists that subsist on a variety of food sources to specialists restricted to only a single host^20–23^. The coupling of host specialization to reproduction raises the possibility that different *Drosophila* species have evolved distinct behavioral adaptations related to the breadth and accessibility of their ecological niches, shaping the structure and dynamics of the social groups that form there. Reproduction and population density in *D. erecta*, for example, are tightly linked to the seasonal availability of its host fruit—the screwpine *Pandanus*^24^—creating a narrow window of opportunity for individuals to meet and mate^25^. By contrast, closely related generalist species, such as *D. melanogaster,* are associated with a variety of perennially available fruits^26^, suggesting that their mating strategies may be adapted to more consistently available environments, where aggregating on a single food substrate is less critical to encounter conspecifics.

Here we show that the olfactory volatiles emanating from a food patch exert both direct and indirect control over mating behavior in *D. erecta*. We find that food volatiles serve as a potent environmental trigger in this species, driving aggregation and serving as a stringent sensory gate to control a male’s sexual arousal. In parallel, environmental food odors shape the visual landscape in which courtship unfolds by encouraging males to approach and sample prospective mates, amplifying visual motion to transform the sensory backdrop of the social scene. *D. erecta* males are highly sensitive to this increase in visual motion, in contrast to *D. melanogaster*, pointing to the evolution of a multisensory feedback loop that coordinates male sexual arousal across members of the group, linking courtship to the broader social context. We show that species-specific differences in the sensitivity to environmental cues appear to arise from variation in how olfactory information impinges on conserved components of courtship circuitry, suggesting a potential neural substrate for the rapid tuning of sensory thresholds that govern sexual arousal. Together, these findings reveal how both ecological and social cues shape species-specific reproductive strategies. Moreover, our observations underscore that the social landscape forms an integral part of the environment—both reflecting the collective dynamics of the group and actively shaping the sensory experience of each individual within it.

## *D. erecta* courtship depends on the olfactory environment

To investigate how environmental factors shape the reproductive decisions of species with different ecological preferences, we compared the dynamics of groups of *D. erecta* and *D. melanogaster* (five males and five females) provided with a small patch of agar perfumed with food-related volatiles (diameter: 21.5 mm) at the center of a large arena (diameter: 190 mm), a configuration intended to mimic the presence of a localized feeding site in the wild (**Fig. 1a**). Although *D. erecta* and *D. melanogaster* have distinct hosts and divergent olfactory preferences^26–28^, both feed on the yeast microbiota that metabolize decaying fruits and thus display attraction to volatiles arising from fermentation^20,22,29^. We therefore used apple cider vinegar (ACV), a broadly appetitive fermentation product as a generic ‘host’ odor to compare how the chemical environment influences mating behaviors. Flies were fed and maintained as virgins in these assays to minimize their motivation to track towards the odor source for purely metabolic needs or in search of an oviposition site.

**Figure 1.**
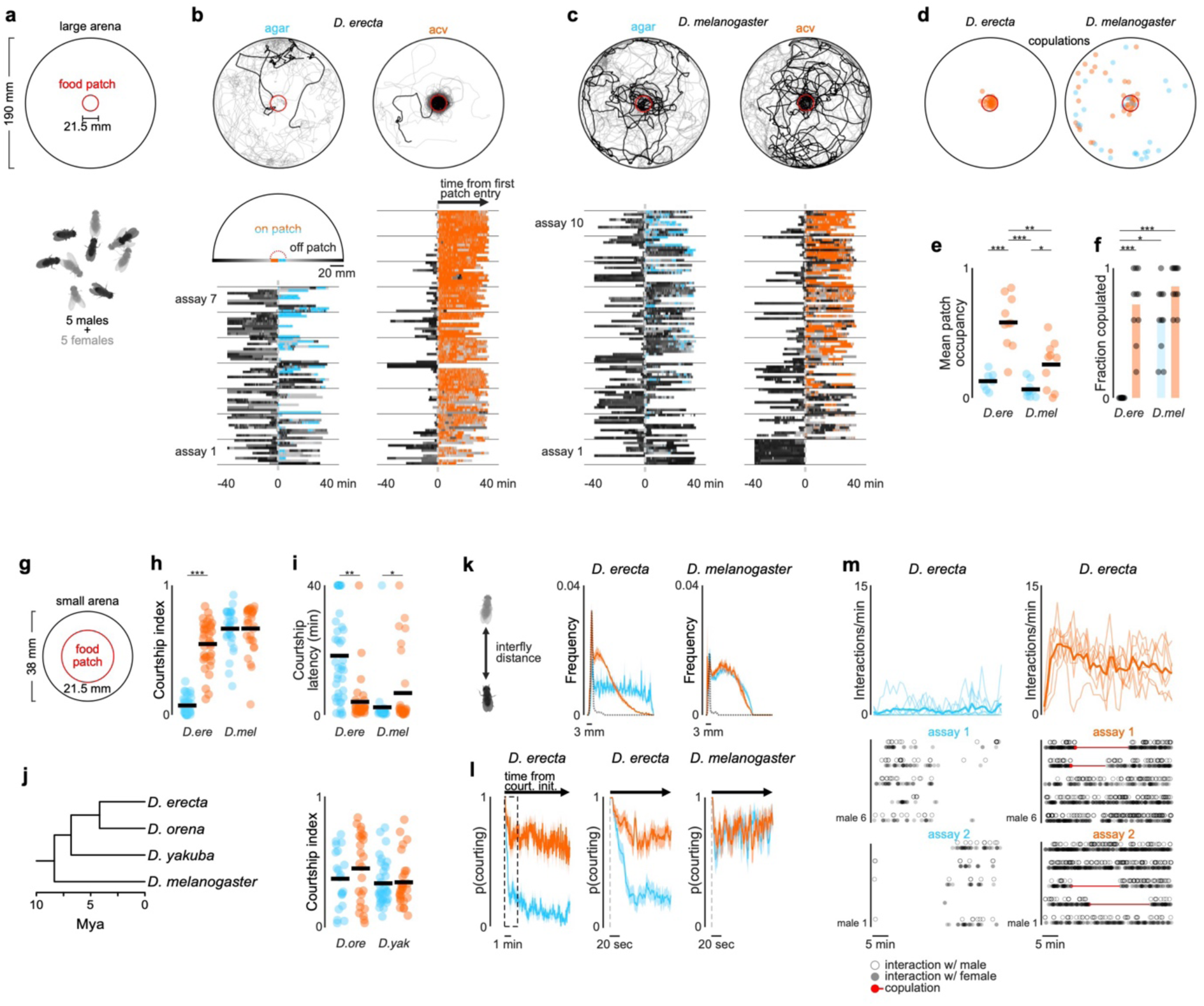
*D. erecta* courtship and mating relies on the presence of ACV. **a**) Schematic of group assay comprised of 5 males and 5 females in large 190 mm arena with a 21.5 mm central food patch formed from agar alone or agar perfumed with ACV. **b**) and **c**) Top: Overlaid trajectories (grey) of representative assays of *D. erecta* (b) and *D. melanogaster* (c) on either agar (left) or ACV (right) with the trajectory of one individual highlighted in black. Bottom: Raster plots depicting the position of each individual fly relative to the agar or ACV patch over duration of the assay (n=7-10 assays). Time on the agar or ACV patch is highlighted in blue and orange, respectively. Greyscale indicates linear distance to the patch with decreasing distance shaded in lighter shades of grey. Individuals are aligned by the time of their first entry to the patch. **d**) Comparison of the spatial distribution of copulations in the presence of an ACV (orange dots) or agar patch (blue dots) for *D. erecta* (left) and *D. melanogaster* (right) (n=10 assays for each species and each condition). **e**) Mean patch occupancy for all flies for *D. erecta* (*D. ere*) and *D. melanogaster* (*D. mel*) with agar (blue) or ACV (orange) food patch (n=10 assays for each species and each condition; t-test with Bonferroni correction). **f**) Fraction of 5 females mated in each 40 min assay for *D. erecta* (*D. ere*) and *D. melanogaster* (*D. mel*) with agar (blue) or ACV (orange) food patch. Each dot represents a separate assay (n=10) and bars represent the mean. **g**) Schematic of group assay comprised of 5 males and 5 females in small 38 mm arena with a 21.5 mm central food patch formed from agar alone or agar perfumed with ACV to examine social interactions at high-density analyzed in (h)-(m). **h**) and **i**) Courtship index (h) and latency to courtship initiation (i) for *D. erecta* (*D. ere*) and *D. melanogaster* (*D. mel*) males (n=6-8 assays for each species and each condition). **j**) Phylogenetic relationships^69^ of *D. erecta* to three closely related species in the melanogaster subgroup (left) and courtship indices of *D. orena* (*D. ore*) and *D. yakuba* (*D. yak*) males in small arena assays (n=4-7 assays). **k**) Distributions of *D. erecta* (left) and *D. melanogaster* (right) inter-fly distances between males and females for the duration of the entire assay (solid line represents mean ± std. err) or just during bouts of courtship (dotted line, mean). **l**) Probability of males courting (mean ± std. err) in the first 10 minutes (left) for *D. erecta* on ACV (orange) or agar (blue). Expanded time window over 1 minute from courtship initiation for *D. erecta* (center, corresponding to dotted box on left) or *D. melanogaster* (right). **m**) Mean interaction rate of all 5 *D. erecta* males towards any other individual in the arena over the duration of an assay (top, solid line represents mean and shaded line individual assays) and ethogram of each male in two representative assays for either agar (blue) or ACV (orange) showing individual interactions with other males (open circle) and females (closed circle). Time of copulation events depicted as red circle (initiation) and line (duration). Dots in (e) and (f) are assay means; dots in (h) – (j) are individual males; horizontal bars in dot plots represent the mean. Those comparisons that were statistically significant are indicated as: * *p* ≤ 0.05; ** *p* < 0.001; *** *p* < 0.0001. *D. ere*: *D. erecta*, *D. mel*: *D. melanogaster*, *D. ore*: *D. orena*, *D. yak*: *D. yakuba*.

We found that in the context of the localized ACV patch, these two ecologically disparate species displayed distinct patterns of behavior. *D. melanogaster* broadly explored the arena, giving rise to a dispersed spatial distribution where individuals interacted mostly in pairs and only intermittently in larger groups (**Fig. 1c**, **Extended Data Fig. 1a-e**). In contrast, *D. erecta* individuals rapidly approached the central ACV patch and remained there for extended bouts (**Fig. 1b,e**; **Extended Data Fig. 1a-e**), forming dynamic clusters of three or more interacting flies. *D. erecta* did not congregate on a patch composed of agar alone (**Fig. 1b,e**; **Extended Data Fig. 1b,f-g**), indicating that food volatiles act as a potent aggregation cue in this species. Consistent with the notion that this aggregation serves a social function in *D. erecta*, over 90% of their copulations occurred on the ACV patch, in comparison to the spatially dispersed mating observed in *D. melanogaster* (**Fig. 1d,f**; **Extended Data Fig. 1h**). Notably, *D. erecta* refrained from mating entirely in the absence of ACV (**Fig. 1f**), highlighting the importance of the food patch as a site of reproductive opportunities in this species.

The ACV patch could facilitate mating in *D. erecta* by simply aggregating individuals, creating a dense social landscape where interactions between males and females are enhanced by proximity. Alternatively, ACV may directly elevate the sexual arousal of males or females upon their arrival to the patch. To differentiate between these possibilities, we assessed the courtship dynamics of groups of five males and five females confined to a smaller chamber (diameter of 38 mm, < 4% of the large arena; **Fig. 1g**), allowing us to decouple the effect of spatial proximity from the influence of the chemosensory environment. *D. melanogaster* males courted vigorously in this arena, independent of the presence of ACV (**Fig. 1h**). By contrast, *D. erecta* males courted only when ACV was present but rarely in its absence, despite the high density of flies (**Fig. 1h-i**). Indeed, *D. erecta* remained dispersed on an agar-only patch (**Fig. 1k**; **Extended Data Fig. 1i**), with males rarely approaching or pursuing individuals of either sex (**Fig. 1m; Extended Data Fig. 2**). In the presence of ACV, however, the spatial distribution of *D. erecta* individuals was transformed, displaying a pronounced peak at 3 mm (**Fig. 1k**) —the inter-fly distance characteristic of courtship pursuit—along with a broad tail, reflecting the dynamic interactions of males as they indiscriminately advanced towards any individual who came into their field of view (**Fig. 1m**; **Extended Data Fig. 2c**). The rare instances of *D. erecta* courtship on agar were transient, in contrast to the persistent pursuit observed in the presence of ACV (**Fig. 1l,m**). ACV thus appears to influence mating behaviors in *D. erecta*, not simply by aggregating males and females but by restructuring their social dynamics.

To investigate whether the influence of ACV was more broadly conserved across the *D. melanogaster* subgroup, we extended our analyses to *D. orena*, a sister species of *D. erecta* that breeds on *Syzygium* waterberries^30^, and *D. yakuba*, a more distant generalist^26^. Both species displayed comparably vigorous courtship in the absence or presence of ACV (**Fig. 1j**), underscoring that *D. erecta* males have evolved a unique dependence on ACV despite the shared ancestry and diverse ecologies of these close relatives.

## Olfactory cues promote social exploration

How does the chemosensory environment transform the structure of the *D. erecta* social landscape? Since interactions between individuals were more frequent on ACV, we reasoned that food odors may heighten the salience of visual objects. To explore this possibility, we presented individual males with a stationary fly-sized target—a high contrast hemispherical bead, 3 mm in diameter (**Fig. 2a**)—and assessed whether ACV altered their interactions with it. *D. melanogaster* males were largely indifferent to this visual target, independent of the presence of ACV, spending most of their time circling the perimeter of the arena. *D. erecta* males, by contrast, remained in close proximity to the bead, an effect that was enhanced by ACV (**Fig. 2b,c; Extended Data Fig. 3a-c,f**). Optogenetic activation of most olfactory sensory neurons using the *Orco* promoter (**Fig. 2d**), which replicates the attractive valence of ACV^31,32^, was sufficient to promote attraction to the bead in *D. erecta* males on agar, confirming that olfactory input underlies this shift in visual salience **(Fig. 2e-f; Extended Data Fig. 3d-e**). Visual approach was not enhanced by the same optogenetic manipulation in *D. melanogaster* males, underscoring that in *D. erecta*, olfactory input exerts a species-specific influence on how males interact with their visual surroundings.

**Figure 2.**
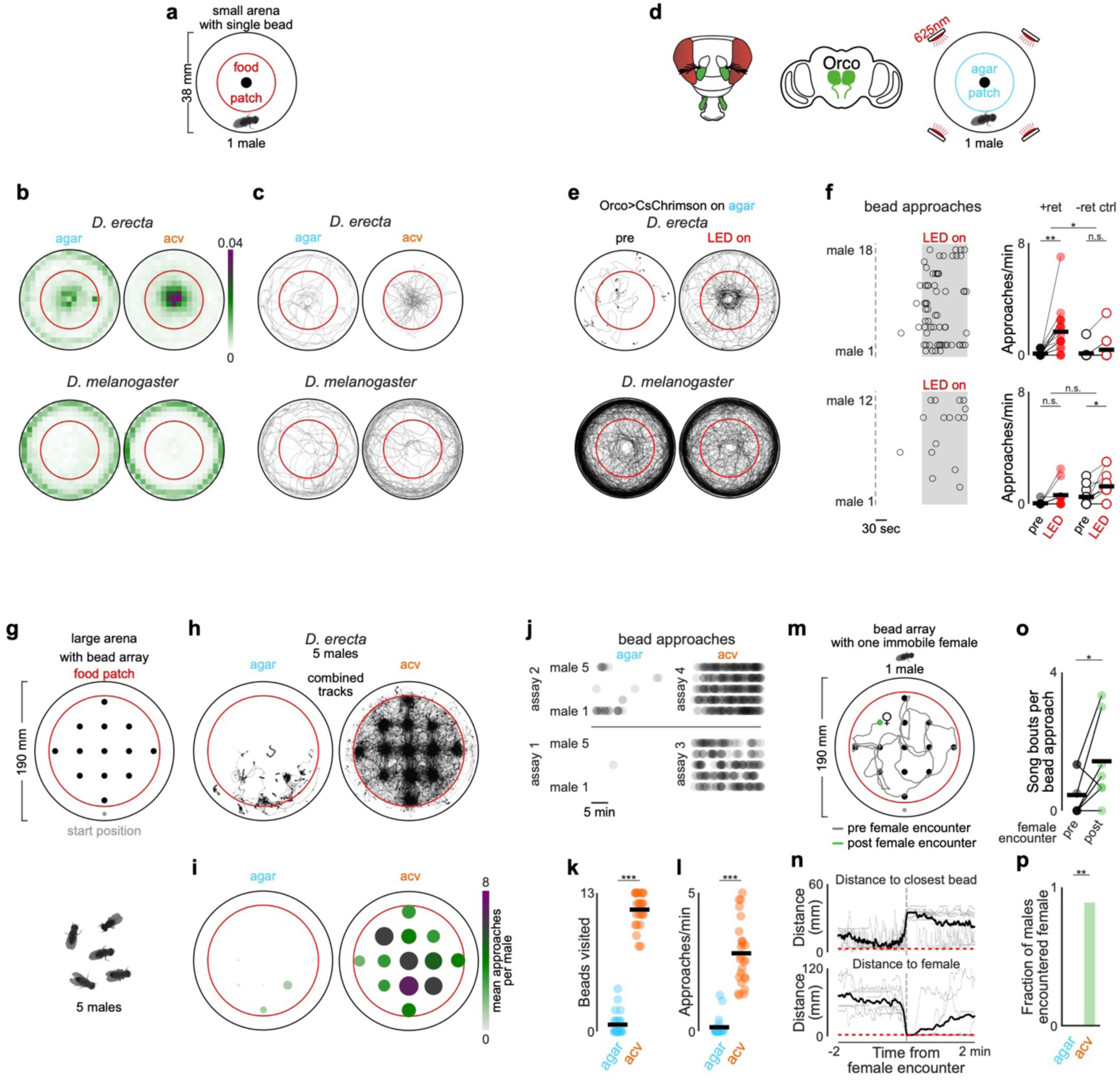
Olfactory cues promote visual exploration in *D. erecta* males. **a**) Schematic of bead assay comprised of a single male with a female-sized hemispherical bead (⌀ ca. 3 mm) at the center of a small 38 mm arena with a 21.5 mm central food patch formed from agar perfumed with ACV or an agar control. **b**) Mean relative distribution of individual *D. erecta* (top, n=15-18) and *D. melanogaster* (bottom, n=8) males on agar (left) and ACV (right). Scale bar indicates relative density. **c**) Overlaid trajectories of males in (b) during the first minute after a fly is within 3 mm of the bead. **d**) Schematic of fly’s head highlighting olfactory appendages (third antennal segments and maxillary palps) innervated by Orco+ olfactory sensory neurons in green and innervation of antennal lobes in the brain (center). Right: Schematic of optogenetic bead assay comprised of a single male with a female-sized hemispherical bead (⌀ ca. 3 mm) at the center in a small 38 mm arena with a 21.5 mm central agar patch. Arena is fitted with LEDs emitting 625 nm (red) light from top. **e**) Overlaid trajectories of *D. erecta* (top, n=18) and *D. melanogaster* (bottom, n=12) Orco>CsChrimson males over 2 minutes prior to (left, pre) and during 2-minute (right, LED on) optogenetic activation. **f**) Left: Raster of individual bead approaches (circles) for each male in (e) in the 2 minutes prior to and during optogenetic activation (grey shaded area). Right: Approaches per minute for Orco>CsChrimson males prior to (black circles) and during optogenetic activation (red circles) for males reared on retinal food (+ret, left, filled circles) and control males of the same genotype but reared without retinal (-ret ctrl, right, open circles). *D. erecta* above and *D. melanogaster* below. Note that light alone modestly enhances approach, independent of rearing condition or resulting optogenetic activation of sensory neurons. **g**) Schematic of grid assay containing a lattice of 13 fly-sized hemispherical visual targets (⌀ ca. 3 mm) arranged in a regular square array in a large 190 mm arena with a large 145 mm food patch formed from agar perfumed with ACV or an agar control. Bead sizes are enlarged and not to scale to highlight grid pattern. **h**) Overlaid trajectories of groups of 5 *D. erecta* males on agar (left) and ACV (right) in the presence of the bead grid (n=5 assays) over 20 minutes. Note that on agar, flies frequently remain close to the position where they were introduced into the arena. **i**) Mean number of approaches per male in (h) for each bead in the grid over 20 minutes indicated by color and size of the circle. **j**) Four representative assays from (h) for agar (left) and ACV (right), in which bead approaches for each male (rows) are depicted (black dots) over the 20-minute assay. **k**) and **l**) Total number of beads in the grid visited (k) and approaches per minute (l) for each individual male in (h) on agar (blue) and ACV (orange). **m**) Schematic of bead grid assay (12 beads, ⌀ ca. 3 mm each) in a large 190 mm arena on a large 145 mm ACV patch with the first bead in the second row from the top replaced with an immobilized female (highlighted with ♀). Example trajectory of a single male colored by time before (grey) and after female encounter (green). Trajectory spans 20 min. Note that after the male encounters the female, trajectory is tightly centered around her. **n**) Distance of males on ACV to the closest bead (above) and the immobilized female (below) centered around the first time the male encounters the female. Thick black line indicates the mean of all males that encountered the female (8 of 9 males), thin lines individuals. **o**) Average number of song bouts as a male approached a bead prior to (black) and post (green) encountering the female for males in (n). **p**) Fraction of *D. erecta* males that encountered the female during the 20-minute assay on agar and ACV. Dots in dot plots represent individual males; horizontal bars in dot plots represent the mean. Those comparisons that were statistically significant are indicated as: * *p* ≤ 0.05; \*\**p* < 0.001; *** *p* < 0.0001.

Despite the strong attraction of *D. erecta* males to the bead on ACV, they frequently walked away before returning to it, generating a distinctive radial pattern as males alternated between approach and retreat (**Fig. 2c**), suggesting that the olfactory environment may also promote visual exploration. To examine this, we placed five *D. erecta* males in a large arena (diameter 190 mm) decorated with a lattice of 13 beads arranged in a regular square array (**Fig. 2g**, distance along edge, 30 mm). In the presence of ACV, males spent most of their time near one of the targets but also frequently traversed between them (**Extended Data Fig. 3g-h**), often taking the shortest route (**Fig. 2h-l**). On agar, males restricted their exploration to a small fraction of the arena, remaining near the bead closest to their starting location and rarely visiting any other visual targets (**Fig. 2h, Extended Data Fig. 3g**). Comparing the trajectories of solitary males revealed that they display a similar shift in their behavior in the presence of ACV (**Extended Data Fig. 3i**), indicating that exploration is driven by the chemical, not social, context. Without the bead lattice, males on ACV showed no spatial bias and remained broadly distributed across the arena (**Extended Data Fig. 3g**). Olfactory cues thus promote exploration in *D. erecta* males.

Beads, while attractive visual targets to males, rarely evoked courtship displays such as the wing extensions associated with song production, suggesting they are not perceived as potential mates (**Extended Data Fig. 3j**). Indeed, replacing a single bead in the array with a decapitated female shifted the search dynamics of males: upon encountering the female, males often terminated their exploration and maintained her at close range (**Fig. 2m-n; Extended Data Fig. 3k**). Males occasionally resumed exploration of nearby beads after encountering the female but now often sang as they approached (**Fig. 2o**), indicating that the distinct cuticular pheromones carried by *D. erecta* females altered the males’ interpretation of visual targets, triggering a shift from exploration to courtship. Males on agar failed to locate the female within the bead lattice (**Fig. 2p**), suggesting that their limited exploration in the absence of food volatiles hinders their ability to detect or pursue prospective female mates. Thus, ACV promotes locomotion in *D. erecta* males to drive more frequent sampling of visual objects in their environment, supporting a form of ‘social foraging’ in which they search for reproductive opportunities rather than food resources.

## ACV promotes male sexual arousal

In *D. melanogaster*, courtship is triggered by recognition of an appropriate conspecific female, a behavioral switch mediated by male-specific P1 neurons that integrate multisensory signals—including direct excitatory input from pheromone pathways—to gate entry into a persistent arousal state^33–37^. *D. erecta* females, like their *D. melanogaster* counterparts, produce sex- and species-specific cuticular hydrocarbons^38,39^ that robustly excite the P1 neurons of conspecific males^40^, suggesting they similarly serve as chemical determinants of mate recognition. Food volatiles could therefore promote *D. erecta* courtship by simply encouraging males to approach and sample female pheromonal cues. Yet even in the rare instances when *D. erecta* males initiated courtship on agar their pursuit was fleeting (**Fig. 1l,m**), pointing to the possibility that food volatiles are required to sustain arousal in this species.

Consistent with this, while optogenetic activation of P1 neurons in *D. melanogaster* is sufficient to evoke persistent courtship^41,42^, in *D. erecta* it triggered only transient courtship bouts towards an immobilized female in the absence of ACV, with males rapidly resuming indifference after the optogenetic stimulus (**Fig. 3a-c**). By contrast, P1 neuron activation drove sustained courtship in *D. erecta* males in the presence of ACV, supporting that while P1 neurons promote a state of sexual arousal, olfactory cues are required for its perpetuation. This olfactory dependence extended to pursuit of a moving visual target: in the absence of ACV, P1 neuron activation evoked only brief bouts of courtship toward a fly-sized 3 mm projected dot rotating on the floor of the assay chamber^43^, whereas in the presence of ACV, males pursued and sang to the target spontaneously, even without P1 activation (**Fig. 3d-h**).

**Figure 3.**
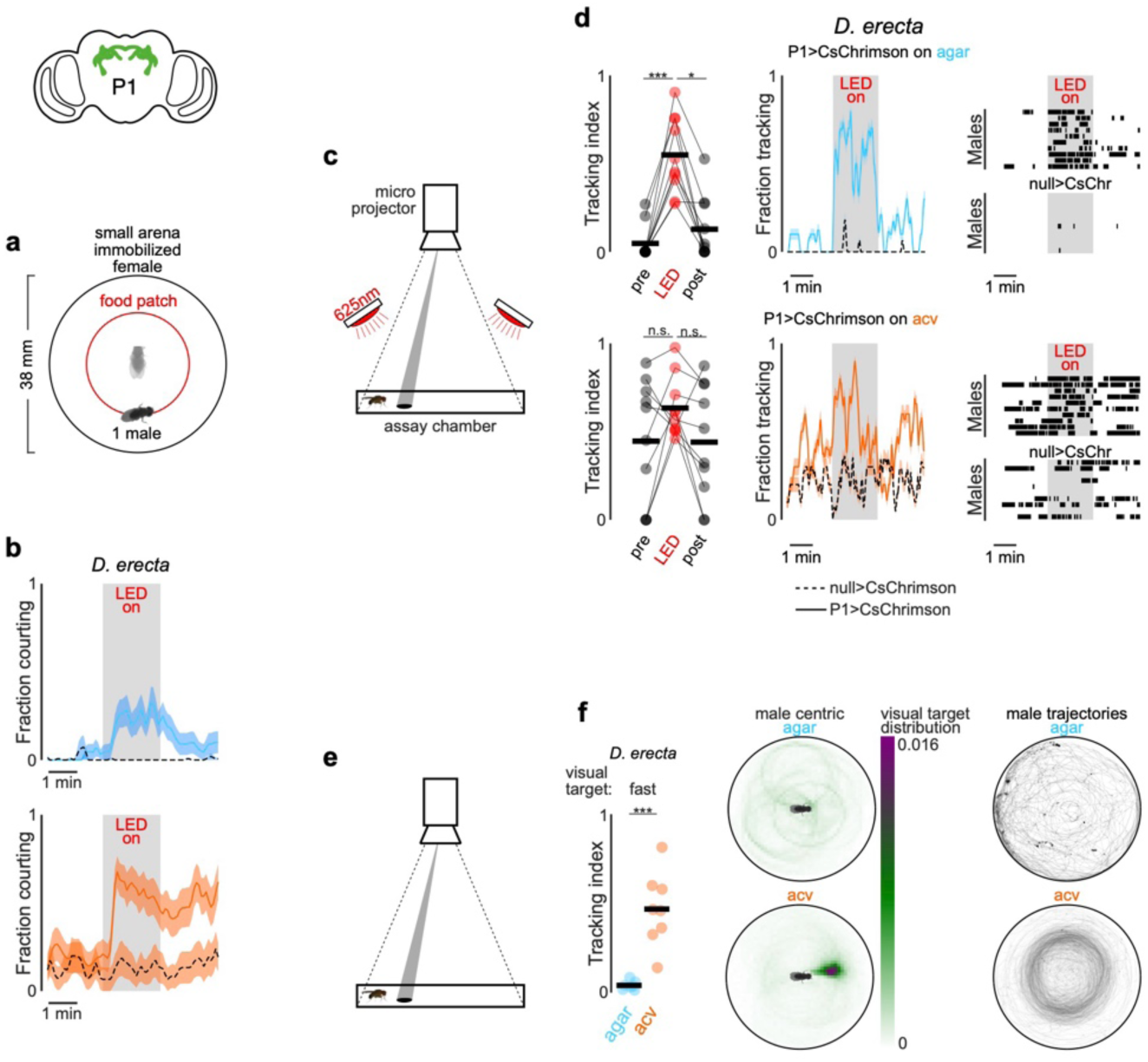
ACV promotes sustained courtship pursuit in *D. erecta* males. **a)** Schematic of optogenetic assay in which a single male is presented with an immobilized (decapitated) female target at the center of a small 38 mm arena with a central 21.5 mm patch made of agar and perfumed with either ACV or agar as control. **b)** Fraction of *D. erecta* males courting the female over time, with optogenetic activation shown as grey bar on agar (above) and ACV (below) for males expressing CsChrimson in P1 neurons (solid line) and genetic controls carrying >CsChrimson transgene but no P1 driver (dotted line). Lines indicate mean ± std. err (shaded) colored in blue for agar and orange for ACV. Note males only have a brief (2 min) period to explore the arena before optogenetic stimulation, in comparison to Fig. 2m. **c**) Schematic of optogenetic projector assay, in which a single male is placed in a small 38 mm arena with either 30 μl agar alone or agar perfumed with ACV in the center and a fly-sized 2-dimensional dot rotating at a constant velocity is projected onto the floor. Arena is fitted with LEDs emitting 625 nm (red) light from the top. **d**) Mean tracking index prior to (pre), during (LED) and after (post) P1 neuron activation (left), fraction of males tracking (center) and raster plot depicting bouts of tracking over time (right) for *D. erecta* males on agar (above) or ACV (below) (n=10 each). Grey box indicates time of LED illumination. Lines indicate mean ± std. err (shaded) colored in blue for agar and orange for ACV. Genetic controls carrying >CsChrimson allele but lacking a P1 driver (null>CsChr) are indicated by black dotted lines (center) and as raster plots (right, bottom). **e**) Schematic of projector assay in which a single wild-type male is placed in a small 38 mm arena with either 30 μl agar alone or agar perfumed with ACV in the center and tracking of a projected fly-sized dot rotating at a constant velocity. **f**) ACV promotes pursuit of a constantly rotating visual target. Left: Tracking index of *D. erecta* males towards a rotating dot in the presence of agar (blue) or ACV (orange) (n=7-8 males). Middle: Heatmap depicting the position of the visual target with respect to the egocentric orientation of the male (facing towards right). Right: overlaid trajectories of males during presentation of the target. n.s.: not significant; * *p* ≤ 0.05; *** *p* < 0.0001.

Food volatiles thus appear to directly regulate the perpetuation of a male’s sexual arousal to orchestrate courtship pursuit in *D. erecta*. To assess whether maintenance of arousal requires continuous olfactory input, we leveraged the fact that tethered males will spontaneously court a fly-sized visual target projected onto a panorama, faithfully tracking its motion while singing to it^42^ (**Fig. 4b**). *D. erecta* males were largely indifferent to this visual target but initiated robust pursuit upon presentation of ACV (**Fig. 4c-f**, **Extended Data Fig. 4**). Notably, courtship tracking was transient, with males terminating pursuit at the offset of the odor stimulus, reinforcing that ACV is required for the perpetuation of arousal (**Fig. 4e-f**, **Extended Data Fig. 4b-c**). *D. melanogaster* males, by contrast, commenced courtship pursuit at the onset of the visual stimulus and persistently tracked it for the duration of the assay (**Fig. 4d-f**), aligned with evidence that visual cues are sufficient to arouse males of this species, even in the absence of additional sensory feedback^37,42^ (**Extended Data Fig. 5**).

**Figure 4.**
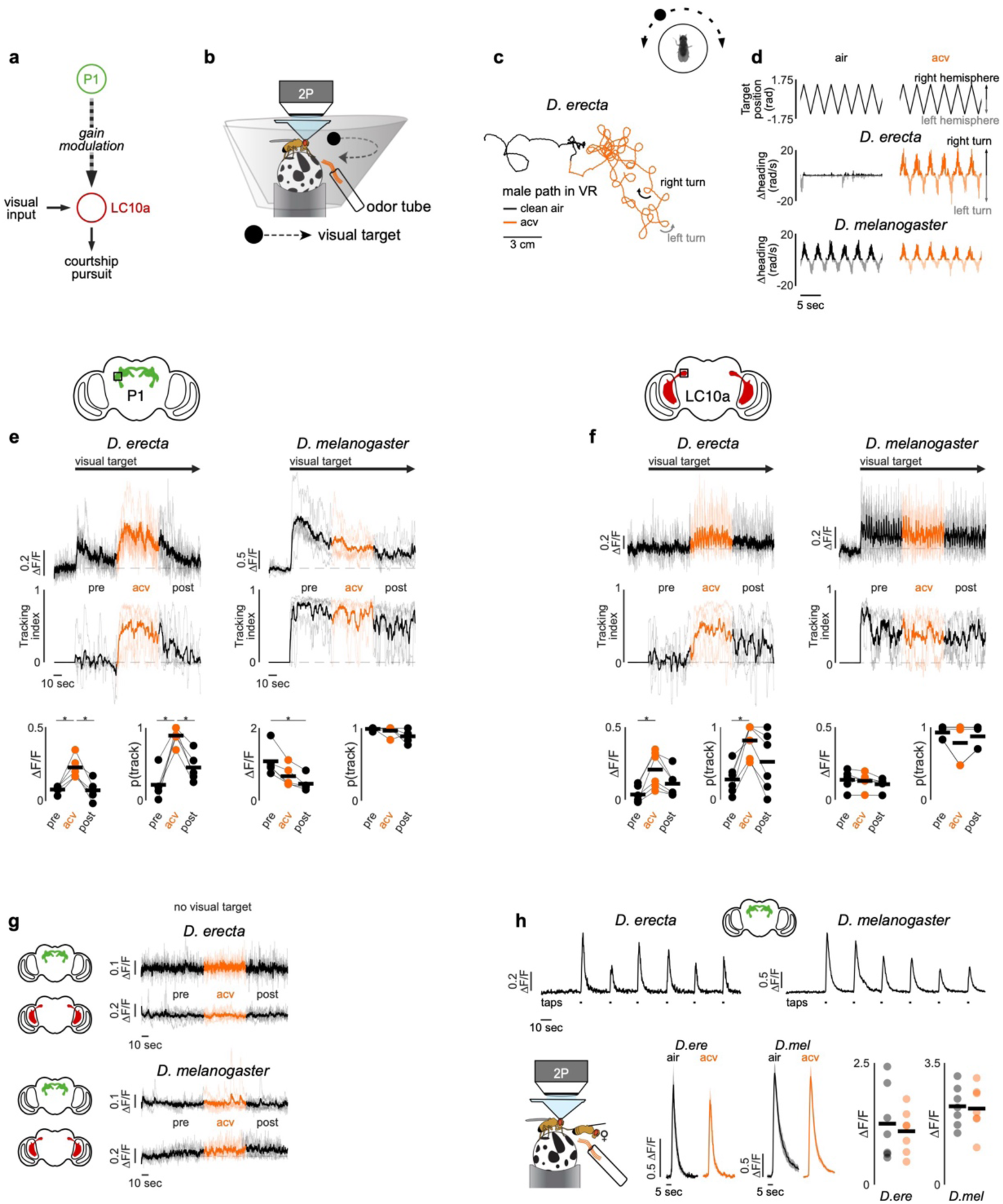
ACV odor recruits central courtship circuitry for persistent pursuit. **a)** Diagram depicting how the gain of LC10a visual projection neurons that mediate courtship pursuit are modulated by P1 neuron activity after Ribeiro et al. (2018)^45^ and Hindmarsh Sten et al. (2021)^42^. **b**) Schematic showing 2-photon imaging setup for tethered flies provided with avisual target (black dot) projected onto a visual panorama rotating at a constant angular velocity and with either clean air or ACV presented through odor tube. **c**) Representative example of *D. erecta* male path in virtual space prior to (black line) and during (orange line) ACV presentation during epochs when visual target is present. **d**) Position of the visual target (top) and corresponding heading direction of an example *D. erecta* (center) and *D. melanogaster* (bottom) male in the absence (left) and presence of ACV (right). **e**) and **f**) P1 (e) and LC10a (f) neural activity (top) and tracking index (center) of *D. erecta* (left) and *D. melanogaster* (right) males prior to (black), during (orange), and after (black) presentation of ACV with visual target continuously oscillating after an initial 30 second baseline period (n= 5-6). Neural activity represented as normalized fluorescence (ΔF/F). Thick lines represent the mean across individual males (thin lines). Arrow indicates appearance of rotating visual target. Bottom: Mean P1 or LC10a neural activity and probability of tracking. **g)** P1 and LC10a neural activity prior to (black), during (orange), and after (black) ACV presentation in the absence of a visual target in *D. erecta* (above) and *D. melanogaster* (below) males. **h**) Top: Representative traces of P1 activity evoked by tapping a conspecific female in either *D. erecta* or *D. melanogaster* males in the absence of odor (clean air). Tick marks represent taps. Below: Schematic of tapping assay (left), mean tap-evoked P1 responses in clean air and ACV odor (center), and corresponding peak responses (right) for *D. erecta* and *D. melanogaster* males. Those comparisons that were statistically significant are indicated. * *p* ≤ 0.05

In *D. melanogaster*, P1 neurons dynamically modulate the gain of LC10a visual projection neurons via a multisynaptic pathway^42,44^ (**Fig. 4a**), forming a concise visuomotor circuit that mediates a male’s faithful courtship pursuit when he is aroused. To assess whether this gain control mechanism is conserved, we generated selective genetic drivers that allowed us to record activity from P1 or LC10a neurons in *D. erecta* males as they engaged in courtship pursuit. Despite the divergent sensory dependencies of courtship across species, the activity of P1 neurons were correlated with bouts of pursuit in both *D. erecta* and *D. melanogaster* (**Fig. 4e**, **Extended Data Fig. 6**). Likewise, in both species, LC10a visual responses were amplified during courtship, with robust activation each time the target swept in the progressive direction across the male’s ipsilateral visual field during epochs of active pursuit (**Fig. 4f**, **Extended Data Fig. 7**). These observations point to a conserved circuit organization in which the gain of LC10a visual responses is dynamically modulated by a male’s sexual arousal to orchestrate courtship pursuit in both species. However, whereas visual cues alone are sufficient to vigorously arouse *D. melanogaster* males, *D. erecta* males additionally require continuous olfactory input, which appears to act as a stringent species-specific gate to sustain courtship pursuit in this species.

Notably, neither P1 nor LC10a neurons were activated by ACV (**Fig. 4g**), indicating that the dependency on food volatiles does not arise from olfactory input to these conserved circuit nodes. Moreover, P1 pheromonal responses remained unchanged by ACV in either *D. erecta* or *D. melanogaster* males (**Fig. 4g-h**), indicating that food odors do not tune the excitability of this integrative node to control a male’s arousal. Rather, ACV activates an array of distributed neurons labeled by the sexually-dimorphic Fruitless and Doublesex transcription factors (**Extended Data Fig. 8**). These include subsets of neurons innervating the anterior optic tubercle to which LC10a send their axonal projections and the lateral protocerebral complex which is innervated by P1 neurons, suggesting a potential substrate through which olfactory modulation acts as a gate via other regulatory nodes downstream of P1 neurons that shape male arousal and sustain pursuit.

## Visual motion promotes courtship

How do environmental food volatiles and visual cues coordinately shape male behavior in naturalistic ecological contexts, where many flies aggregate on a food patch? By driving *D. erecta* males to sample and pursue visual targets (**Fig. 2**), ACV results in an increase in fly locomotion, not apparent at the level of individual pairs, but rather that scales with the size of the group (**Extended Data Fig. 9a,c**). In the context of larger aggregates of flies, this odor-dependent shift dramatically altered each male’s visual experience (**Fig. 5a-e**)—expanding the fraction of his visual field occupied by other flies (**Fig. 5d**) and increasing the visual motion sweeping across his retina (**Fig. 5a-c,e**). The motion generated by interacting individuals on a food patch thus transforms the visual backdrop, suggesting it may convey information about social context to control male arousal. Consistent with this, even on agar, males displayed correlated bursts of locomotor activity and courtship (**Fig. 5f-h, Extended Data Fig. 9b**), indicating that motion cues emanating from nearby flies transiently elevate male arousal and pointing to a feedback loop in which ACV both promotes courtship and reshapes the sensory landscape in which courtship unfolds.

**Figure 5.**
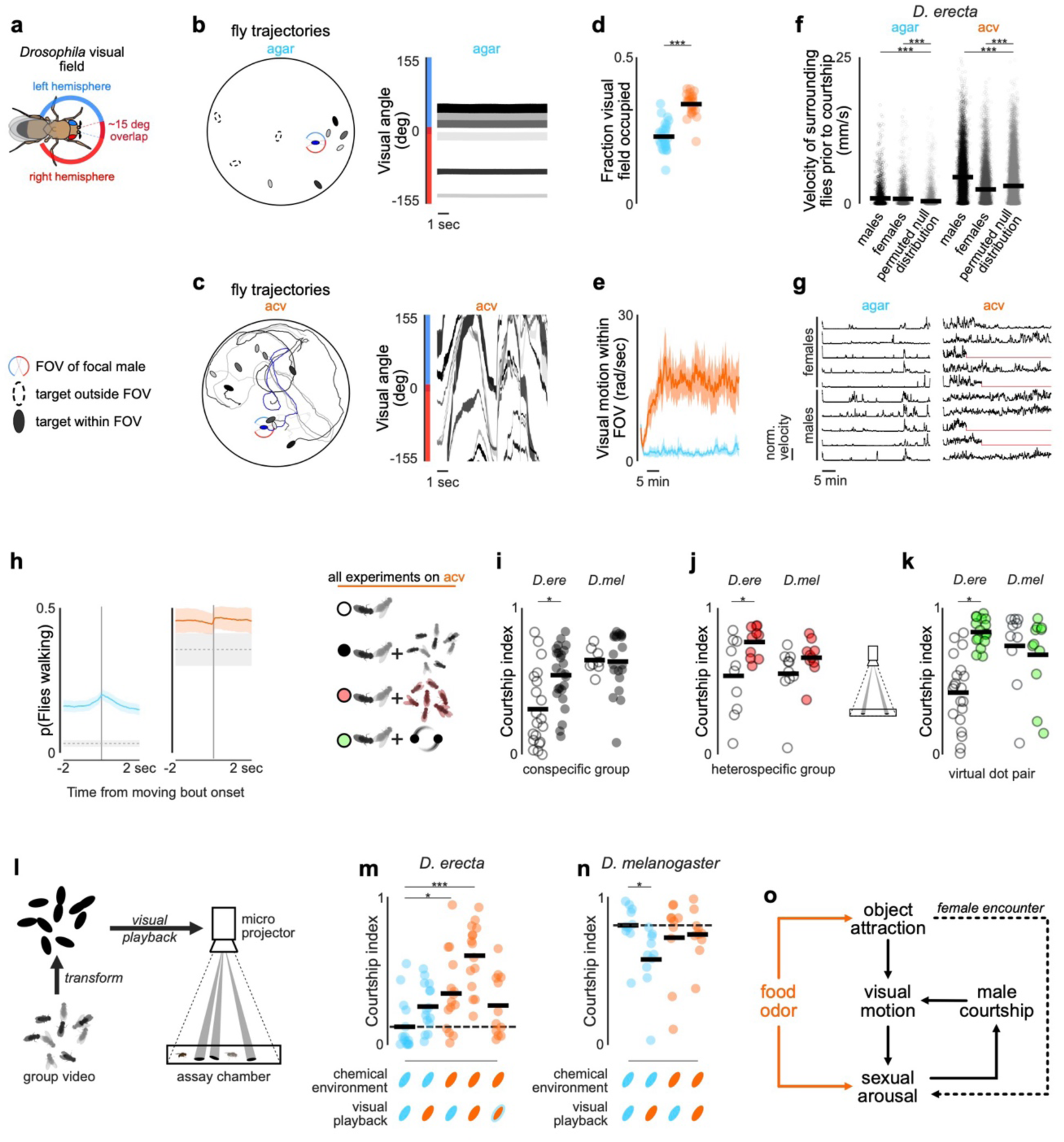
Visual motion cues modulate *D. erecta* courtship in groups. **a**) Schematic of the *Drosophila* visual field across the left (blue) and right (red) eye and indicated ∼15 degrees of binocular overlap. **b**) and **c**) Left: Representative trajectories and final positions of all individuals within a group of 5 males and 5 females over a 10 second period. Flies were in a 38 mm arena with a 21.5 mm central food patch composed of agar (b) or agar perfumed with ACV (c). Individuals within the field of view (FOV) of a focal male (blue oval) are shown as filled and those outside his FOV are shown as dotted ovals in greyscale. Paths of individuals shown as different shades of grey. Right: Projection of other individuals within the group over the 10 second period onto the focal *D. erecta* male’s visual field. Colors of individual trajectories match greyscale on the left. **d**) and **e**) Mean fraction of the visual field of *D. erecta* male occupied by other individuals (d) and mean visual motion within the visual field of view (FOV) (e) throughout the 40-minute recordings of 5 males and 5 females in a 38 mm arena with a central 21.5 mm patch filled with agar perfumed with ACV (orange) or an agar control (blue). **f**) Mean velocity of all other males and females in the group during the 2 second window preceding a male courtship bout, compared to a permuted null distribution of 2-second velocity windows in *D. erecta* groups on agar (left) and ACV (right). **g**) Representative example depicting velocity normalized by maximum velocity per individual for all flies over the duration of a 40-minute assay composed of 5 males and 5 females in a 38 mm arena with a central 21.5 mm patch filled with agar (blue) or agar perfumed with ACV (orange). **h**) Probability of other flies walking given movement bout onset for any individual fly within D. erecta groups (n=7-8 assays) on agar (left, blue) and ACV (right, orange) compared to randomly shuffled data (1000 replicates per assay) highlighted in grey. Means and standard error across assays are indicated by thick lines and shaded areas, respectively. **i**) and **j**) Courtship indices of *D. erecta* (*D. ere*) and *D. melanogaster* (*D. mel*) males when in groups of 5 males and 5 females (filled circles) or paired with just one female (open circles) in small 38 mm arenas with a central 21.5 mm patch formed from agar perfumed with ACV. Males in groups were surrounded by either conspecific group (i, black) or a heterospecific group of 4 male and 4 female *D. simulans* (j, red). **k**) Courtship indices of *D. erecta* (*D. ere*) and *D. melanogaster* (*D. mel*) males paired with a single conspecific female in a 38 mm arena with 30 ul agar perfumed with ACV in the center (open circles) in comparison to the same condition where two fly-sized projected dots rotated at constant angular velocity (one slow (16 mm/sec) and one fast (32 mm/sec) target) on the floor of the arena (closed circles, green). **l**) Schematic of visual replay experiment. Individual trajectories recorded from a group of 5 males and 5 females in a small 38 mm arena with a central 21.5 mm ACV or agar patch were tracked and used to create representative videos of the distinct group dynamics in each chemosensory environment by approximating each individual’s centroid position as a black ellipse. These videos were replayed to a test male and female pair by projecting them onto the floor of a 38 mm arena to assess how male courtship behavior was modulated by either replay of the visual scene (recorded from groups of flies on ACV or agar) or the current chemical environment (when ACV or agar control present in the testing arena). **m**) and **n**) Courtship indices of males towards a conspecific female for all combinations of chemical environment and visual replay for *D. erecta* (m) and *D. melanogaster* (n). Flies were either placed in an arena containing an agar control (blue ellipse) or ACV (orange ellipse) and paired with visual playback recorded from a group on agar (blue ellipse) or ACV (orange ellipse) or the ACV video slowed to the average speed of flies in the agar video (orange ellipse with blue outline). The courtship indices of each individual male are shown with circles colored by the respective chemical environment (agar: blue, ACV: orange). **o**) Diagram depicting the direct and indirect effect of food odor on the sexual arousal and social behaviors of *D. erecta* males within an aggregate of conspecifics. Comparisons that were statistically significant are indicated as: * *p* ≤ 0.05; *** *p* < 0.0001.

Aligned with the synergistic effect of food odors and visual motion on arousal, we found that individual *D. erecta* males on ACV displayed more vigorous courtship when part of a larger group (five pairs of conspecifics) than when presented with just a single conspecific female in isolation—an effect not observed in *D. melanogaster* (**Fig. 5i**). Courtship towards a conspecific female was similarly enhanced by the presence of the same sized group of *D. simulans*, underscoring that these courtship promoting cues are not species-specific. Indeed, male courtship in *D. erecta* was similarly elevated by simply projecting two continuously rotating dots onto the assay chamber floor, highlighting that visual motion is sufficient to promote arousal (**Fig. 5j-k**).

Male courtship in *D. erecta* thus appears to be influenced by both the chemosensory environment of a food patch as well as the dynamic motion of the social group on it. To explore the composite effects of food odor and visual motion, we recorded the group dynamics of five pairs of *D. erecta* males and females on agar or ACV, approximated each fly’s centroid position as a high contrast ellipse, and then played back these visual motion cues to a ‘test’ *D. erect*a male and female pair (**Fig. 5l**). We observed courtship in *D. erecta* males was enhanced by playback of the visual scene from flies interacting on ACV (**Fig. 5m)**. Courtship of the test pair on ACV was comparable when presented with either the agar visual scene or a slowed version of the ACV scene that matched the velocity of flies on agar, suggesting that *D. erecta* males are sensitive to the speed of visual targets, rather than their distinct dynamics. Notably, males on ACV, presented with the visual motion cues recorded on ACV, displayed the highest level of courtship, supporting the synergistic effects of the chemosensory and visual environment. Neither the presence of ACV nor playback of the social scenes recorded on ACV further increased the vigorous courtship displayed by *D. melanogaster* (**Fig. 5n**). Together, our data suggest that the chemosensory environment of a food patch promotes and perpetuates a male’s sexual arousal in *D. erecta* by driving males to sample and pursue visual targets, which in turn reshapes the visual social scene to further reinforce mating behaviors (**Fig. 5o**).

## Discussion

In *Drosophila*, courtship unfolds as a tightly coupled feedback loop between prospective mates, continuously shaped by the reciprocal exchange of sensory signals between male and female. Here, we examined how this feedback loop is influenced by the ecological and social context of *Drosophila* mating sites, revealing that in the host specialist *D. erecta*, both the olfactory volatiles emanating from a food patch and the collective dynamics of flies congregating there coordinately structure the sensory environment to control courtship. We find that food odor regulates male courtship in *D. erecta* through two distinct mechanisms: first, by gating a male’s arousal to promote pursuit of potential mates, and second, by further enhancing his arousal through the increased visual motion of other individuals on the food patch, creating a feedback loop that promotes courtship in environments where reproductive opportunities are most abundant*. D. erecta* has thus evolved a striking dependence on environmental cues as an integral part of its reproductive strategy.

*D. erecta*’s distinct sensitivity to its environment is an adaptation well aligned with its specialization on *Pandanus* fruit, a seasonally restricted host that is only transiently available for reproduction. Breeding populations of *D. erecta* surge during the two-to-four-month period when *Pandanus* fruit is present, then sharply decline as the availability of fruit wanes^24,25^. During seasons when *Pandanus* is scarce, *D. erecta* can opportunistically subsist on other fruits, but at greatly diminished numbers, pointing to the inherent cost inherent of specialization on ephemeral resources. Such strict dependence underscores how *Pandanus* serves as a key ecological hub supporting the reproductive cycle of this species^24,25^.

Despite *D. erecta’s* tight ecological association with *Pandanus*, we find that ACV is sufficient to broadly restructure its mating behaviors. *D. erecta’s* specializaGon may thus reflect long-range akracGon to the disGnct volaGles of its host^28^, whereas once at the fruit, more general chemical signatures of fermentaGon control sexual arousal and maGng. ACV contains many metabolites of fermentation that play a broader role in drosophilid ecology^46,47^. Indeed, the ability of ACV to promote courtship appears to be maintained in a latent form in closely-related *Drosophila* species—emerging only when males are presented with a relatively impoverished two-dimensional visual target (**Extended Data Fig. 5**) or in darkness^48^. Likewise, ACV has been shown to alter the salience of visual objects during flight in *D. melanogaster*^49,50^, indicating that the arousing effect of food volatiles is broadly conserved, with its behavioral expression quantitatively tuned by species-specific sensory thresholds.

In *D. erecta*, the sensory threshold for arousal appears particularly high, rendering courtship strictly dependent on continuous olfactory input. Indeed, transient activation of P1 neurons is insufficient to drive the sustained pursuit that other *Drosophila* species display^41,42^, suggesting that olfactory input acts downstream of P1 neurons to perpetuate a male’s arousal state. Consistent with this idea, ACV not only sustains P1-mediated arousal but can independently promote courtship pursuit. While the precise neural locus of ACV’s action remains to be elucidated, our findings suggest that the gating of courtship by food volatiles in *D. erecta* arises from conserved circuit components that differ quantitatively across species to generate distinct thresholds for arousal. For example, *D. melanogaster* can become highly aroused with just visual input, consistent with evidence that subsets of P1 neurons receive direct LC10a input (Gerry Rubin, personal communication). P1 neuron activity in turn modulates the gain of LC10a responses^42^, forming a recurrent circuit poised to perpetuate visually guided courtship pursuit. One intriguing possibility is that ACV-responsive olfactory pathways gate this feedback loop in *D. erecta,* such that both vision and food volatiles are required for males to sustain their sexual arousal.

Such subtle variation of conserved circuit elements underscores how courtship circuits can diversify while maintaining their core function, enabling the rapid adaptation of different species to their distinct ecological needs. The latent influence of food volatiles in related species, for example, suggests that variation in sensitivity to environmental cues may arise through the differential weighting of existing sensory pathways rather than the recruitment of novel inputs—a principle reminiscent of the diversification of pheromonal tuning in P1 neurons^51^. Pheromone responses in P1 neurons, for example, are unaffected by ACV (**Fig 4h**), indicating that olfactory and pheromonal pathways remain segregated, an organization that preserves the ability of males to discriminate between conspecific and heterospecific females, even when highly aroused on a food patch (**Extended Data Fig 10**). The modularity of courtship circuits^40,52^ thus favors the independent diversification of distinct circuit elements, allowing specialists, such as *D. erecta*, to evolve unique behavioral sensitivity to environmental cues and exploit the potentially fleeting reproductive opportunities their host fruit provides.

Beyond the direct modulation of circuits perpetuating sexual arousal, ACV also serves to structure the social landscape at multiple levels. First, ACV triggers *D. erecta* flies to rapidly track to the odor source. Flies in our experiments were fed and sated, suggesting their aggregation on a food patch is driven by reproductive rather than metabolic needs. Second, once on the food patch, ACV promotes males to sample visual targets, presumably in search of sexually receptive females, transforming the visual scene through their collective motion during this form of ‘social foraging’. Finally, because *D. erecta* males are highly sensitive to visual motion, their courtship is further enhanced by the presence of moving visual targets. This in turn orchestrates the behavior of nearby individuals such that their bouts of pursuit become synchronized, an effect most apparent on agar, where periods of immobility are punctuated by intermittent bursts of activity across the group.

Aggregating in larger groups increases a male’s access to potential mating partners, but at the same time promotes competition, which is thought to be fierce on a crowded food patch^53–55^. Reproductive success in these competitive environments requires that males fend off their rivals and isolate a female for long enough to initiate copulation^56^. As a consequence, a male’s sensitivity to visual motion may confer an advantage in gaining access to potential mating opportunities, by triggering him to chase after any visual object that may be a receptive female or courting pair. Indeed, on a food patch, males become attracted to all visual objects and indiscriminately approach a male, female, or a bead. The drive to compete therefore likely structures the collective activity observed in groups of *D. erecta*.

Although *Drosophila* is not typically thought to engage in collective social behaviors, the dynamics of *D. erecta* on a food patch may follow the same principles that structure schooling fish^57^, swarming locusts^19^, or murmuring starlings^58,59^. In these systems, simple local interaction rules—such as attraction and repulsion among neighbors— align the behavior of many individuals into concerted actions^19,60–62^. Likewise, in *D. erecta* aggregates, the behavior of neighboring flies serves as an environmental reference that modulates courtship behavior—not through direct signaling between prospective sexual partners, but through generic visual motion cues that indicate the broader environmental context. However, in contrast to canonical examples of collective behaviors—where information is thought to be shared to optimize group fitness—*D. erecta’s* sensitivity to group dynamics likely evolved to enhance individual reproductive fitness. Our results thus suggest that collective behaviors can emerge from diverse selective pressures, giving rise to coordinated actions even in species not generally considered ‘social’.

## Material and Methods

### Fly stocks and husbandry

Flies were housed at 25C and 50-65% relative humidity on a 12h light/12h dark cycle, except for animals used during optogenetics experiments, which were kept in darkness. Stocks used for behavioral experiments were reared on standard molasses food, 2P imaging lines were reared on Würzburg food for at least five generations to increase animal robustness, and flies expressing channelrhodopsins were kept on sugar-yeast food to prevent intake of the co-factor retinal. Fly stocks used were as follows: *D. melanogaster* stocks 71G01-Gal4 (39599) and UAS-GCaMP6s (42746) obtained from Bloomington Stock Center, Canton-S (G. Maimon), UAS-CsChrimson (V. Jayaraman), VT029314 (LC10a)-LexA, UAS-CsChrimson, LexAop-GCaMP6s (I. Ribeiro and B. Dickson). Wild type *D. erecta* (14021-0224.01), w^-^ *D. erecta* (14021-0224.07), and *D. yakuba* (14021-0261.00) were obtained from the Cornell Stock Center, Ruta lab *D. simulans*, *D. orena* (D. Stern). *D. erecta* 71G01-Gal4, UAS-GCaMP6s, UAS-CsChrimson, ppk23-mutant (Coleman et al. 2024); Dsx-Gal4 (Y. Ding). *D. erecta* VT029314 (LC10a)-Gal4, Fru-Gal4, Orco-Gal4, ΔOrco and ΔIr8a were generated in this study.

### Transgene generation

Vector backbone pRC12B^40^ with ie1.mCherry minigene reporter was used to clone the synthesized VT029314 enhancer fragment gBlock (IDT) upstream of UAS via Gibson assembly (NEB). The construct was co-injected with hyTsps mRNA into embryos for transposase-mediated genome integration. Chemosensory co-receptor knockouts were generated by targeted excision of the first three exons (Orco) or the entire coding region (Ir8a) using CRISPR/Cas9-mediated homology directed repair inserting a fluorescent marker gene cassette encoding mCherry under the viral ie1 promoter (Orco) or RFP under control of *D. melanogaster* 3xP3 (Ir8a). The Orco-Gal4 line in *D. erecta* was generated by cloning 5496 bp upstream of the Orco translation start site upstream of the Gal4 sequence into the pCaSpeR backbone and a gateway casette (pCaSpeR-DEST3 (DGRC Stock 1029; RRID: DGRC_1029). The fru-Gal4 line was generated by first establishing an attP landing site within the *D. erecta* fruitless locus and the subsequent insertion of a splice acceptor Gal4 element. The landing site was created using CRISPR/Cas9-mediated cleavage to the intron 5’ of the exon containing the ATG start codon following^63^ with homology directed repair inserting the attP sequence. The resulting validated *fru^attP^* line was used as target for site-specific integration of a splice acceptor-Gal4 element, which contained an ie1.mCherry transformation marker.

### Free-behavior assays

All free-walking behavioral assays were conducted at 25C and 50-65% relative humidity between 0h and 3h after lights on. Virgin males and females were collected after eclosion and group housed separated by sex for 4-7 days before the assay. Flies were aspirated without anesthesia, females first followed by males. All free-behavior videos were taken at 30 or 42 frames per second (fps) in back light from a white light pad (Logan Electric) using a Point Grey FLIR Grasshopper camera (GS3-U3-32S4M-C 3MP) fitted with a HF12.5SA-1 lens (Fujinon). Central food patches of varying sizes (see below) were formed either from agar (2% w/v) perfumed with ACV (20% v/v) or a 2% agar-only control.

#### Large Arena

Assays were performed in 190 mm diameter, 3mm high circular slope walled Delrin chambers modeled after the flybowl^64^ with a central food patch of 21.5 mm diameter. Five females were transferred to the chamber by mouth aspiration followed by five males. Once the males were loaded the assay commenced, and groups were recorded for 40 minutes. Single-individual assays lasted 20 minutes. The centroid position of each fly was tracked in every frame and identity swaps were manually corrected using the Caltech flyTracker^65^. The positional data was then used to infer individual trajectories, relative distributions, and the distance of flies to the central food patch in MATLAB (MathWorks). A clustering fly was defined as within 3 fly lengths of at least one other fly, whereas a solitary individual was defined as being at a distance of at least 3 fly lengths of all other flies in the arena. The timing and position of copulations were manually scored.

#### Grid Arena

Grid assays were conducted in modified large 190 mm diameter arenas with a large 145 mm diameter central food patch. A grid forming a lattice of 13 black, fly-sized (3 mm diameter) hemispherical plastic beads (Etsy) arranged in a regular square array was placed on the patch. The grid consisted of 5 equidistant beads along the horizontal midline with 30 mm distance along the edge, 3 equidistant beads 30 mm above and below and one additional bead 30 mm above and below the three beads all aligned along the midline. Assays with either 5 males or a single solitary male were recorded for 20 minutes. For ‘social foraging’ assays, one randomly chosen bead of the grid was exchanged with an immobilized decapitated female. Recording commenced after aspiration of all males. The centroid position and speed of each fly was tracked in every frame using AnimalTA^66^. If necessary, identity swaps were corrected manually within the software. Exact bead positions were extracted for each assay using the same software. Distances of males to beads or the immobilized female and visits to visual targets were calculated from the positional data and used to infer individual trajectories and relative distributions. Summary statistics including the number of beads visited per male, bead approaches over time, the number of times a male switched between any two different beads, and the fraction of time spent per bead or with the female were calculated from positional data in MATLAB. A visit was defined as an individual spending at least 30 seconds within three fly lengths (∼9 mm) of a bead or the female. The timing of an approach was defined as the first frame of a visit bout. Bead switches were defined as the visit of any bead different than the previous one. Male unilateral wing extensions (singing) during the approach of a bead or a female were tracked manually using Boris^67^.

#### Small Arena

Performed in 38 mm diameter, 3 mm high circular slope walled Delrin chambers with a central food patch of 21.5 mm diameter. Videos were recorded in 2 x 2 grids using the same recording setup as above. Females were aspirated first followed by males and assays commenced after the last fly was loaded for a 40-minute recording. For heterospecific group assays, 4 pairs of *D. simulans* females and males were transferred to the assay chamber first, followed by one *D. erecta* female and one male. Ppk23-knockout assays were performed by first transferring a *D. melanogaster* female target to the arena followed by mouth aspiration of either a wild type or a *D. erecta* ppk23 mutant tester male on ACV. In single-bead assays, a hemispherical black plastic bead of ∼3mm diameter (Etsy) was placed on the food patch at the center of the assay chamber first followed by aspiration of a single fly after which the assay commenced. Bead assays were recorded for 20 minutes. The centroid position and orientation of all flies were tracked and manually corrected for identity swaps in flyTracker. Reproductive behaviors of males were then inferred in MATLAB from his orientation, heading direction and position relative to visual stimuli and other flies in the arena. Specifically, we extracted male approach and tracking of other flies, close-range courtship, and copulations. Approach was defined as the directed movement (≥0.8 mm/sec) towards a sedentary object, such as another fly or a plastic bead from within a distance of maximally 4 fly lengths (∼12 mm). We scored approach when the male’s orientation was within 60 degrees of the line connecting his and the target’s center of area up until a distance of 1.5 fly lengths (4.5 mm) where it had to be within 45 degrees. Courtship was defined as periods during which males were approaching or tracking another fly or were within 1.5 fly lengths (∼4.5 mm) of a female with an angle of at least 45 degrees subtending the line connecting the male’s and female’s centroid. We calculated the courtship index as the fraction of time a male was engaged in courtship for the duration of the assay. In case a male copulated, courtship index indicates the fraction of time a male was engaged in courtship until copulation. CI= Time spent courting/Assay length. To calculate the visual experience of males within groups in different chemosensory environments, the angular position and width of other flies in the arena on the focal flies visual field were calculated in MATLAB on a frame-by-frame basis as previously described^44^.

### Free-behavior visual virtual reality (VR) projector assays

All free-behavior visual VR assays were performed in 38 mm diameter, 3 mm high slope-walled chambers made of Delrin, coated with white self-adhesive vinyl paper (Livelynine) used to enhance reflectance increasing contrast of projected shapes. Visual stimuli were directly projected onto the assay chamber surface^43^ using a DLP LightCrafter 4500 mini projector (Texas Instruments). Tester males were aspirated into the assay chamber and allowed to acclimate for 5 min before visual stimulus projection started and recording commenced for 5 minutes using a Basler acA3088-57uc camera with a 8 mm/F 1.8 lens (Edmund optics). If pairs of flies were used, the female was aspirated first followed by the male after which the assay commenced and was recorded for 40 minutes. All videos were recorded at 30 fps. For all assays either 30ul 2% (w/v) agar perfumed with ACV (20% v/v) or a 30ul 2% agar-only control were pipetted into the center of the assay chamber and allowed to solidify before flies were aspirated.

#### Constant angular velocity stimulus assays

Males were presented with either a slow dot rotating clockwise at 16 mm/sec with a 7.5 mm radius from the center of the assay chamber, a fast dot rotating counter clockwise at 32 mm/sec with a 15.5 mm radius from the center of the assay chamber, or both simultaneously. Each dot had a diameter of 3 mm. For mating assays, a female target was added and both visual stimuli were presented after the aspiration of the male when the assay commenced for 40 minutes or until copulation, whichever was first. The positions of the visual stimuli, female, and the male were tracked using flyTracker and male visual object tracking behavior was inferred in MATLAB based on his orientation, heading direction and position relative to the visual stimuli and female. Tracking was defined as the directed movement towards a moving object within the orientation criteria described for approach in free-behavior small arena assays above. The data was then used to calculate the tracking index for each male. The tracking index was defined as the fraction of time a male tracked a visual stimulus during the course of an assay. TI=Time spent tracking/Assay length.

#### Visual environment replay assays

The visual environment replay videos were prepared as follows. Individual trajectories tracked and manually corrected with flyTracker were extracted from one representative recording of a *D. erecta* group of 5 males and 5 females on ACV and one recorded on agar in a small arena. The positional centroid information as well as the orientation of each individual were then used to place a black ellipse at each fly’s position with the long axis (length: 3mm) aligned to the orientation for each frame of a total of five minutes recorded at 30 fps. The five minutes were then looped to create a 40-minute video for each chemosensory condition. This was done, because on ACV the accumulation of copulating pairs over time changes the visual environment of a group from one dominated by dynamic movement during courtship to a more sedentary one. We thus extracted a representative 5-minute block of a group dominated by courtship on ACV. For the agar video we extracted the 5-minute block at the same time point. For the assay, replay of either the agar or ACV visual environment was paired with agar (2% w/v) or agar perfumed with ACV (20% v/v). To not obstruct the video replay, four equidistant 30 ul dollops were placed in a square formation at the periphery of the assay chamber. A female and a male fly were aspirated into the arena and the recording started together with video replay. Male behavior towards the female was scored manually using Boris for 40 minutes or until copulation, whichever came first. Manual tracking was necessary, because flies were obscured at low contrast by the ellipse projections at times, which was not detectable by automated software. The behavioral data was then used to calculate the tracking index for each male.

### Tethered VR assays

#### Tethering

Males (4-5 days old) were briefly (<30 s) anaesthetized on CO^2^, and subsequently tethered to a custom-milled plastic head plate similar to those used in previous studies^42,68^. Flies were held in place by a string across the cervical connective and fixed to the head plate by both eyes and the back of the thorax using UV-curable glue (Loctite 3106 and Bondic). The string was subsequently removed, and flies were left to recover in a warm, humidified chamber (25 °C, 50–70% humidity) in the dark. Flies were transferred to an air-supported ball of 5 mm diameter after 30 minutes and left to acclimatize for 45 minutes.

#### Virtual reality system

We adapted a previous hardware design for presenting tethered males with visual stimuli^42^ (J. Weisman & G. Maimon unpublished) and extended it to provide odors with temporal precision. The system described in detail in Hindmarsh Sten et al. (2021) consisting of an aluminum base keeping an air supplied Styrofoam ball afloat, held in place by a custom 3D printed contraption was adapted to fit either the original 6.35 mm ball used for *D. melanogaster* males or a 5 mm ball for *D. erecta* males. The conical screen surrounding the ball was kept at the original dimensions (large diameter: ∼220 mm, small diameter: ∼40 mm, height: ∼60 mm) but was cut from holographic rear projection film (Megicolim). The visual stimulus was rear projected around the male using a DLP 3010 Light Control Evaluation Module (Texas Instruments). A high-contrast circular visual stimulus of ∼28° diameter mimicking the angular size of a female fly 2 mm away from the male was rear projected onto the screen, moving in a symmetric 150° arc at constant angular velocity (50 °/sec). For odor delivery, we fit Teflon tubing to the setup providing a constant stream of humidified air with low flow volume (0.5 l/min) towards the male’s head from the front at a 45° angle and ∼2 mm distance from the fly as to not obstruct the view on the screen. The tubing was 1 cm in diameter making sure the entire male was engulfed by the stream of air. Pure ACV odor was mixed with water vapor to a concentration of 50% by two digital mass flow controllers and fed into the outlet tubing by switching from pure air using solenoid valves controlled via a data acquisition device (MMC 1408 FS). This ensured air flow rate was constant throughout the experiment. Acquisition of the male’s positional read-out as well as the regulation of visual stimulus generation and odor delivery were combined in MATLAB on a single computer.

#### Experimental parameters

Each assay lasted 210 seconds beginning with a 30 second baseline period followed by the projection of the oscillating visual stimulus for the remaining 180 seconds. ACV odor was delivered for 60 seconds beginning 90 seconds after assay start, exposing the male to 60 seconds of visual stimulus only before and after odor delivery.

#### Tracking index

The tracking index was defined as previously described^42^ to reflect the fidelity and vigor of the male’s pursuit of the visual target.

### Optogenetic assays

Optogenetics experiments were performed expressing the channelrhodopsin CsChrimson in olfactory sensory neurons (Orco-gal4) or P1 neurons (71G01-gal4) with flies reared on sugar-yeast (SY) food lacking retinal, a cofactor required for channelrhodopsin function. After 5 to 6 days, tester flies were transferred to food containing 400 uM all-trans-retinal for two days before the experiment. Controls were either transferred to a new SY food vial or alternatively to food containing retinal when using a genetic control. Genetic control males carried CsChrimson without no Gal4 driver preventing its expression. For optogenetic activation, flies were exposed light at a wavelength of 625 nm emitted by to red LEDs (Cree Xlamp XP-E2). We used three different experimental configurations:

#### Free behavior optogenetic assay

Small 38 mm assay chambers with a central 21.5 mm food patch formed from agar (2% w/v) perfumed with ACV (20% v/v) or an agar control were backlit with blue light. A female, immobilized by decapitation was placed at the center of the arena. After aspiration of the male, assays were run for six minutes, with a with two two-minute blocks of baseline background illumination interleaved by the delivery of red light pulsing at 5 Hz with a pulse length of 0.1 seconds for two minutes. Red light for optogenetic activation was delivered from below the assay chambers using an array of 9 high-power LEDs emitting red light at high intensity to penetrate the Delrin resulting in a power of ∼7uW at the surface. Male behavior was tracked using flyTracker. His position, orientation relative to the female, and heading direction were used to track courtship behavior and the calculate courtship index as described above.

#### Free behavior projector optogenetic assay

Small 38 mm assay chambers were coated with a white adhesive film to increase reflectance and illuminated from above using a LightCrafter 4500 mini projector (Texas Instruments) set to emit light exclusively via its blue LED channel by switching off the red and green LEDs for background lighting. P1 neurons labeled with 71G01-Gal4 were activated by red light delivered from above using 5 high-power LEDs. Assays were run for six minutes, with two two-minute blocks of baseline background illumination interleaved by two-minute delivery of constant red light in addition to the background light. Male behavior was tracked using flyTracker. His position, orientation relative to the female, and heading direction were used to calculate tracking index. Tracking was defined as the directed movement towards a moving object within the orientation criteria described for approach in free-behavior small arena assays above. The data was then used to calculate the tracking index for each male. The tracking index was defined as the fraction of time a male tracked a visual stimulus during the course of an assay. TI=Time spent tracking/Assay length. Population behavior was summarized by the fraction of males courting across assays over time.

#### Tethered optogenetics assay

Males used for optogenetics assays were tethered as described above in ‘Tethered VR Assays’. Assays lasted 210 seconds beginning with a 30 second baseline followed by visual stimulus delivery for the remaining 180 seconds. One 1.5-mm optic fibre (Edmund Optics) was coupled to a high-power red LED (650 nm, LED Engin) mounted on a heat-sink (Ohmite) and placed directly above the head of the fly. Orco+ olfactory sensory neurons were activated by pulsing light at 5 Hz with a pulse length of 0.1 seconds for 60 seconds after 90 seconds. Tracking index was calculated as previously described^42^.

### Two-photon functional imaging

#### Fly preparation

Males were tethered as described above. After recovery, the plate was filled with saline (108 mM NaCl, 5 mM KCl, 2 mM CaCl_2_, 8.2 mM MgCl_2_, 4 mM NaHCO_3_, 1 mM NaH_2_PO_4_, 5 mM trehalose, 10 mM sucrose, 5 mM HEPES, pH 7.5 with osmolarity adjusted to 265 mOsm) covering the fly’s head. The cuticula above the antenna and in-between the eyes was cut with a 30-gauge needle and removed with forceps without anesthesia to provide optical access to the central brain. The trachea covering the top of the brain were removed using forceps after which flies were transferred to the ball to recover in darkness for 45 minutes.

#### Functional imaging

Experiments were performed with an Ultima two-photon laser scanning microscope (Bruker Nanosystems) with a Chameleon Ultra II Ti:Sapphire laser. Samples were excited at a wavelength of 920 nm and emitted fluorescence was detected with a GaAsP photodiode detector (Hamamatsu). Images were acquired with a 40x Olympus water-immersion objective with 0.8 NA. To reduce noise caused by light emitted by the projector during imaging, the red and green LED channels were switched off. In addition, a piece of blue-light filter paper (Rosco) was placed in front of the projector lens. After the male’s resting period on the ball, the objective was lowered over the brain to identify the brain region of interest. We centered a small ROI yielding an imaging rate of 4-8 Hz. Laser power was kept low to reduce stress on the fly. ROIs were drawn in a single hemisphere dependent on the neural population imaged. For P1 and LC10a imaging, an ROI was drawn in the LPC and AOTu neuropil, respectively, where axonal projections were densest. For *fru* and *dsx* imaging, a larger ROI was drawn, encompassing neurites in both the LPC and adjacent AOTu to capture neural activity simultaneously. Experimental parameters and calculation of the tracking index as described in ‘Tethered VR Assays’.

#### Imaging analysis

Imaging data was post processed and analyzed using FIJI (NIH 2.14.0). For each experimental recording, frames were motion corrected and aligned along the x and y axes by the Template Matching plugin. Residual background noise from the projector was reduced from raw time series using the *Despeckle* function. Finally, an ROI was drawn across the entire population of interest containing neuropil and the average fluorescence extracted. Fluorescence was normalized in MATLAB for each recording using the average fluorescence over the first 30 seconds of an experiment as baseline (F_0_). ΔF/F was defined as ΔF_i_/F_0_=(F_i_-F_0_)/F_0_, where i denotes the current frame. Because imaging data were collected at lower frame rate than behavioral data, behavioral data was downsampled using linear interpolation at the imaging time points.

### Statistical analysis

MATLAB was used for statistical analyses and plotting. The Shapiro-Wilk method was used to test normality, and the appropriate parametric/non-parametric statistics was used. In cases for which multiple comparisons were made, appropriate post hoc tests were conducted as indicated in Extended Data Tab. 1. All statistical tests used were two-tailed.

## Acknowledgements

We thank S. R. Datta, D. Kronauer, Y. Ding, J. Rhee, A. Ryba and C. Dowell for comments on the manuscript; G. Maimon, V. Jayaraman, I. Ribeiro, B. Dickson, Y. Ding, and D. Stern for sharing fly lines; J. Weisman and G. Maimon for sharing designs for their projector-based visual VR system presenting tethered flies with a visual stimulus in advance of publication; M. Hathiyari, T. Hindmarsh Sten, R. Li, P. Stock, P. Strogies, J. Petrillo, and Rockefeller Precision Instrumentation Technologies for technical advice; N. Parker for sharing his grid assay design; L. Vosshall for access to the 2-photon system; R. Tollino for assistance in generating the Orco.gal4 line; and the members of the Ruta lab for discussion. Stocks obtained from the Bloomington Drosophila Stock Center, NIH P40OD018537, the Cornell NDSCC Stock Center, and FlyBase were used in this study. A vector obtained from the Drosophila Genomics Resource Center, NIH 2P40OD010949, was used in this study. This work was supported by the Simons Foundation (718234), a Kavli Neural Systems Institute Fellowship and NIH NIGMS grant K99GM151471 (PB); an NIH NIGMS Medical Scientist Training Program grant T32GM152349 to the Will Cornell/Rockefeller/Sloan Kettering Tri-Institutional MD-PhD Program (NBE); a Helen Hay Whitney Foundation Fellowship and NIH NIGMS grant K99GM141319 (RTC); an ERC Starting Grant (802531), and by the Francis Crick Institute, which receives its core funding from Cancer Research UK (CC2067), the UK Medical Research Council (CC2067) and the Wellcome Trust (CC2067) (LP-G); an NIH NINDS grant R35NS111611 (VR) and the Simons Foundation Collaboration for the Global Brain (VR). VR is an Investigator of the Howard Hughes Medical Institute.

## Author contributions

PB and VR conceived of and designed this study. PB designed and performed experiments, analyzed data, and interpreted the results, with input from VR. PB designed and generated transgenic lines with input from RTC. KK carried out grid assay and Δppk23 experiments, contributed to large arena assays and assisted with the generation of transgenes. RTC designed and generated the Fru-gal4 driver line in *D. erecta*. NBE conducted visual environment replay experiment. SZ assisted with initial courtship assays and cloning. LP-G designed and generated the Orco-gal4 driver line in *D. erecta*. PB and VR wrote the manuscript with input from all authors.

## Correspondence

Correspondence should be addressed to Vanessa Ruta.

## Competing Interests

The authors declare no competing interests.

## Code and data availability

Code and data underlying this study are available upon reasonable request from the corresponding author.

---

**Extended Data Figure 1.**
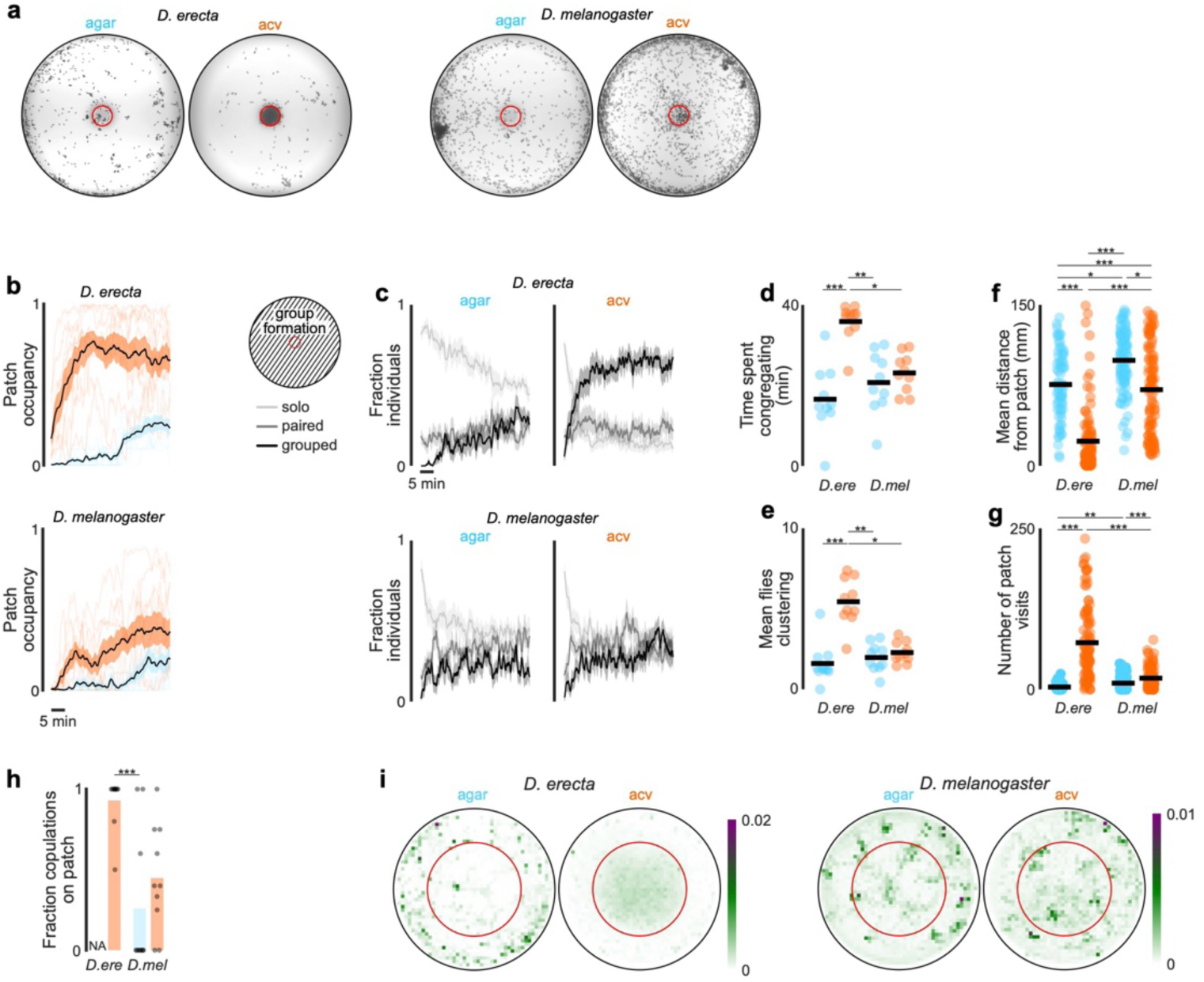
*D. erecta* aggregates on a central food patch in the presence of ACV. **a)** Projections of the distribution of every fly over the first 15 minutes of representative 40-minute group assays with 5 males and 5 females in a large 190 mm arena with a 21.5 mm central food patch (red circle) formed from agar perfumed with ACV (orange) or an agar control (blue) for *D. erecta* (left) and *D. melanogaster* (right). The positions of each fly in every 250th frame are overlaid, projecting a total of 108 frames over 15 minutes. **b)** Population occupancy of the central patch over time for 40-minute group assays with 5 males and 5 females in a large 190 mm arena with a 21.5 mm central food patch for *D. erecta* (above) and *D. melanogaster* (below). Occupancy ranges from 0 with no individuals on the patch to 1 when all 10 individuals of the group are located on the central patch at a given time. Means ± standard error are indicated in black and shaded regions for assays on an ACV (orange) or agar patch (blue). Patch occupancies of individual assays are shown in shaded lines (n=10 each). **c)** Fraction of individuals of group assays in (b) within 3 fly body lengths of one other fly (paired, black), in a group of three or more flies (grouped, dark grey), or at a distance over 3 body lengths from any other individual (solo, light grey) over time for *D. erecta* (above) and *D. melanogaster* (below). All flies in the entire arena were included in the analysis. Means ± standard errors are indicated. **d)** Total amount of time at least one group of three or more flies is present in assays with 5 males and 5 females in a large 190 mm arena with a 21.5 mm central food patch formed from agar perfumed with ACV (orange) or an agar control (blue) for *D. erecta* (D.ere) and *D. melanogaster* (D. mel). Means (black bars) and individual assays (dots, n=10 each) are shown. **e)** Mean number of flies in large arena (190 mm) group assays (5 males and 5 females) present in clusters of three or more individuals throughout the whole 40-minute assay for *D. erecta* (D. ere) and *D. melanogaster* (D. mel) in the presence of an agar (blue) or ACV (orange) patch. Assay means (dots; n=10 each) and overall mean (black bar) are shown. **f)** and **g)** Mean distance of individuals in large 190mm arena from the central 21.5 mm food patch formed from agar and perfumed with ACV (orange) or an agar control (blue) (f) and number of patch visits (g) for *D. erecta* (D. ere) and *D. melanogaster* (D. mel) over 40 minutes. 100 flies from a total of 10 assay each (dots) and overall means (black bar) are shown. **h)** Fraction of all copulations per large 190 mm arena group assay (5 males and 5 females; n=10 assays each) that commenced on the central 21.5 mm patch formed from agar perfumed with ACV (orange) or an agar control (blue) for *D. erecta* (D. ere) and *D. melanogaster* (D. mel). **i)** Mean relative distribution of flies in small 38 mm arena group assays with 5 males and 5 females in the presence of a central 21.5 mm food patch (red circle) formed from agar perfumed with ACV (right) or an agar control (left) for *D. erecta* (left panel; n=7-8) and *D. melanogaster* (right panel; n=5-6). Scale bar indicates relative density. Comparisons that were statistically significant are indicated as: * *p* ≤ 0.05; \*\**p* < 0.001; *** *p* < 0.0001

**Extended Data Figure 2.**
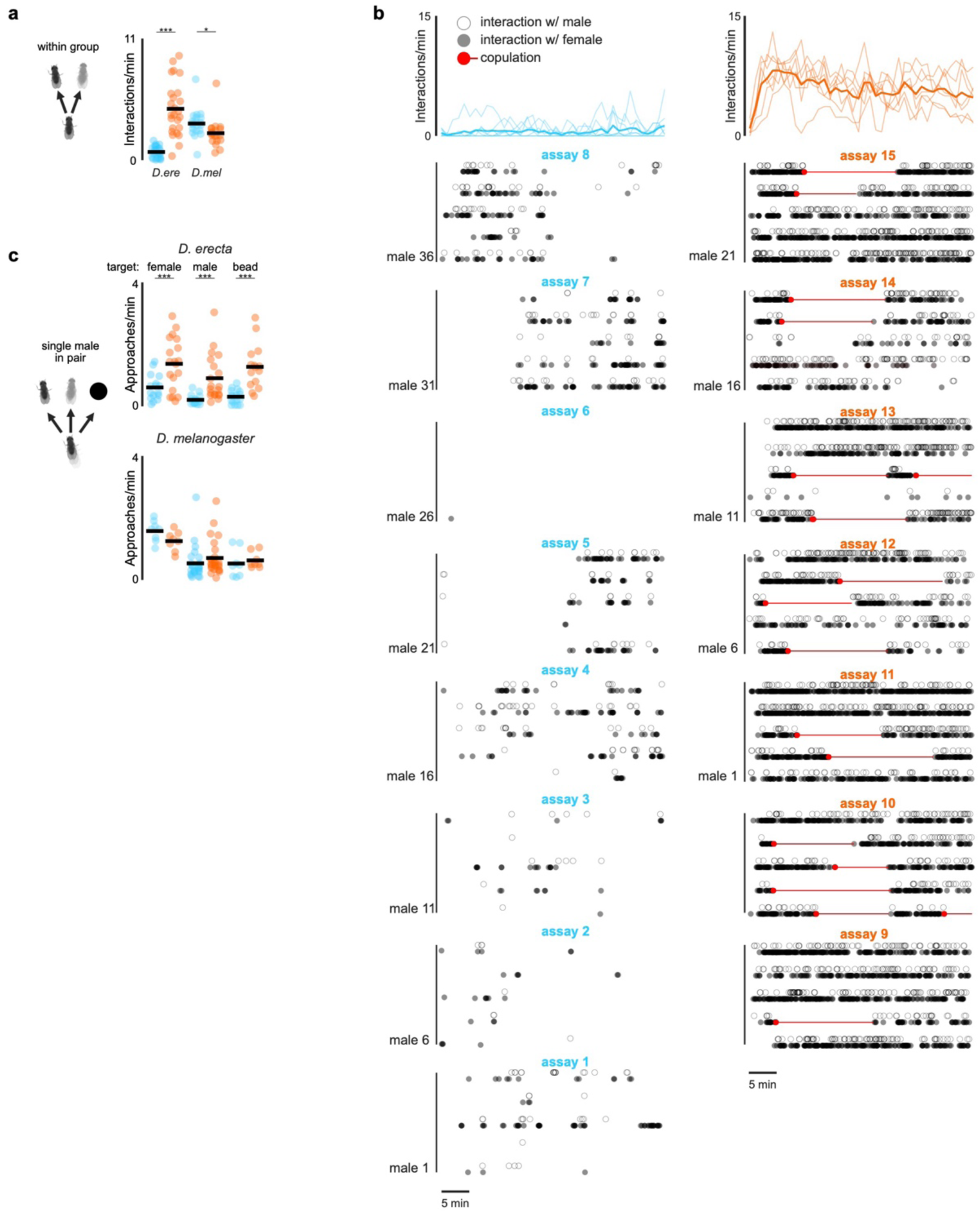
ACV promotes approach of other flies and visual objects in *D. erecta* males. **a)** Interaction per minute of all five *D. erecta* males of 5 male and 5 female group assays in small 38 mm arenas with a central 21.5 mm food patch towards any other individual in the arena for *D. erecta* (D. ere) and *D. melanogaster* (D. mel). Individual males (dots) and means (black bars) are indicated for assays with ACV (orange) or an agar (blue) patch. **b)** Interactions of all five males towards any other individual in the arena over the duration of an assay (top, solid line represents mean and shaded line individual assays) and ethogram of each male for either agar (blue; n=8) or ACV (orange; n=7) showing individual interactions with other males (open circle) and females (closed circle). **c)** Approaches per minute of individual males towards a single female, another male, or an inanimate fly-sized hemispherical visual target (⌀ ca. 3 mm) in a small 38 mm arena with a central ACV (orange) or agar (blue) patch for *D. erecta* (top; n=10-19) and *D. melanogaster* (bottom; n=8-12). Means (black bar) for all assays (dots) are indicated. All males were paired with a single mobile target (another male or female) or a single hemispherical bead positioned at the center of the arena. Those comparisons that were statistically significant are indicated as: * *p* ≤ 0.05; *** *p* < 0.0001.

**Extended Data Figure 3.**
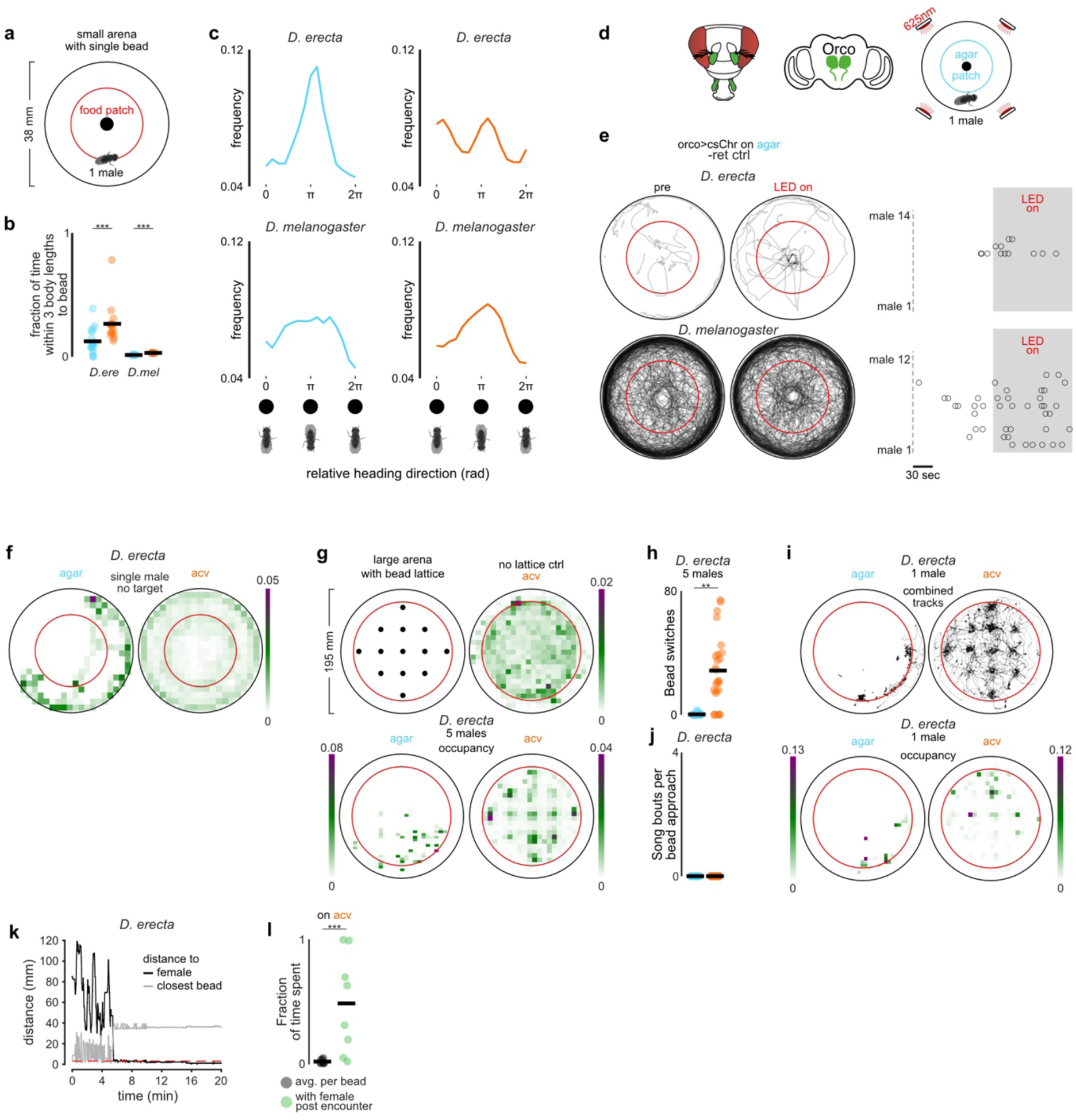
ACV promotes exploration of visual objects in *D. erecta* males. **a)** Schematic depicting assay in which a single male was provided a female-sized hemispherical bead (⌀ ca. 3 mm) at the center of a small 38 mm arena with a 21.5 mm central food patch (red circle) formed from agar perfumed with ACV or an agar control. **b)** Fraction of time individual *D. erecta* (*D. ere*, left, n=15-18) and *D. melanogaster* (*D. mel*, right, n=8) males (dots) spent within three body-lengths of the hemispherical bead at the center of a small arena in the presence of an ACV (orange) or agar (blue) patch over 20 minutes. Mean is indicated by black bars. **c)** Frequency distribution of the central hemispherical bead position relative to the heading direction of *D. erecta* (top, n=15-18) and *D. melanogaster* (bottom, n=8) males in (b) on an agar (blue, left) or an ACV (orange, right) patch. Coordinates are shown using orientations of all males over 20 minutes with respect to the egocentric orientation of the male, heading towards the bead at 0/2π, while the bead is behind the male at π. **d)** Schematic of a fly head highlighting olfactory appendages (third antennal segments and maxillary palps) innervated by Orco+ olfactory sensory neurons in green and innervation of antennal lobes in the brain (center). Right: Schematic of optogenetics bead assay comprised of a single male with a female-sized hemispherical bead (⌀ ca. 3 mm) at the center in a small 38 mm arena with a 21.5 mm central agar patch. Arena is fitted with LEDs emitting 625 nm (red) light from top. **e)** Left: Overlaid trajectories of *D. erecta* (top, n=14) and *D. melanogaster* (bottom, n=12) Orco>CsChrimson control males reared without retinal over the 2 minutes prior to (left, pre) and during 2-minute (right, LED on) optogenetic activation. Right: Raster of individual bead approaches (circles) for each male in the 2 minutes prior to and during optogenetic activation (grey shaded area). **f)** Mean distribution of males in 38 mm arenas with one isolated male in the presence of a central 21.5 mm food patch (red circle) formed from agar perfumed with ACV (right) or an agar control (left) for *D. erecta*. No visual target was provided in these assays. Scale bar indicates relative density. **g)** Top left: Schematic of grid assay containing a lattice of 13 fly-sized hemispherical visual targets (⌀ ca. 3 mm) arranged in a regular square array in a large 190 mm arena on a large 145 mm food patch (red circle) formed from agar perfumed with ACV or an agar control. Bead sizes are enlarged and not to scale to highlight grid pattern. Top right: Mean distribution of groups of 5 *D. erecta* males on ACV without lattice of hemispherical visual targets over 20 minutes. Bottom: Mean distribution of groups of 5 *D. erecta* males in the presence of lattice on an agar (left) or ACV (right) patch. Scale bar indicates relative density over 20 minutes. **h)** Number of times any single *D. erecta* male within the group of 5 males switched from one hemispherical visual bead target to another on an agar (blue, left) or an ACV (orange, right) patch (n=5 assays, 5 males each) over 20 minutes. **i)** Top: Overlaid trajectories of individual *D. erecta* males on agar (left) and ACV (right) in the presence of the bead grid (n=10 assays) over 20 minutes. Bottom: Mean distribution of individual *D. erecta* males on an agar (left) or ACV (right) patch in the presence of a bead lattice. Scale bar indicates relative density. **j)** The number of song bouts per bead approached for *D. erecta* males in grid assay containing a lattice of 13 fly-sized hemispherical visual targets in a large 190 mm arena on a large 145 mm food patch formed from agar perfumed with ACV (orange) or an agar control (blue). Individual males were alone in the assay chamber and behavior was scored for all approaches during 20 minute assay. **k)** Distance of a male to the immobilized (decapitated female,black) and the closest bead in representative grid assay shown in Fig. 2m with the bead in the second row on the left replaced by an immobilized decapitated female over 20 minutes. Red dotted line indicates one fly length (3 mm) distance. Once male has encountered the female, he remains close to her for the remainder of the assay, moving farther from the rest of the beads. **l**) Average fraction of time a male spent with each bead prior to encountering the female (black dots) and fraction of time male spent with the female after encountering her (green dots) for all males that encountered the female on ACV (8 out of 9). Those comparisons that were statistically significant are indicated as: \*\**p* < 0.001; *** *p* < 0.0001.

**Extended Data Figure 4.**
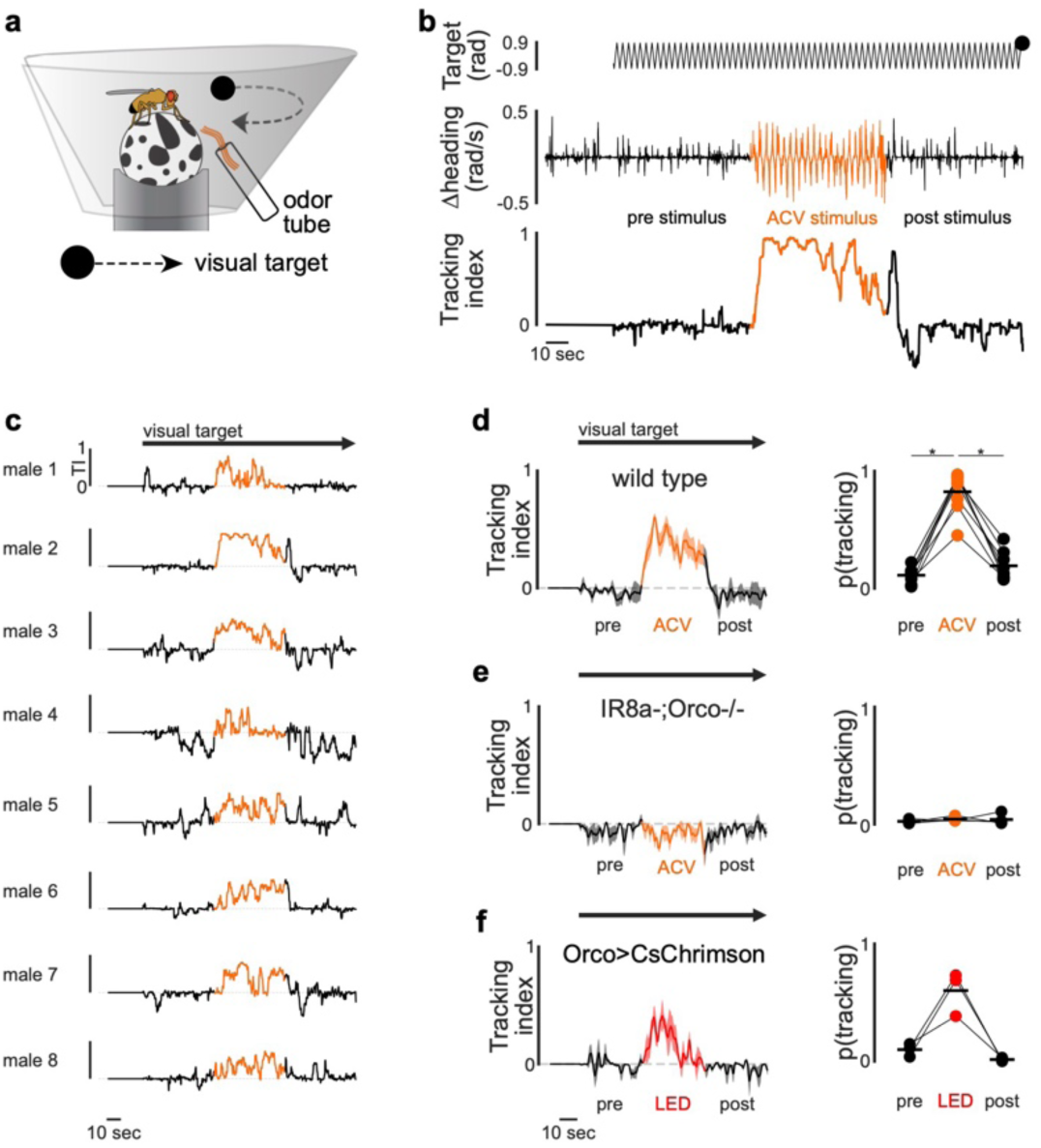
Stimulation of olfactory pathways triggers courtship tracking. **a)** Schematic showing virtual reality behavioral setup for tethered flies provided with an oscillating visual target (black dot) and with ACV presented through odor tube. **b)** Target position in the male visual field (top) and corresponding heading direction (center) and tracking index (bottom) of a representative *D. erecta* male prior to (black), during (orange), and after (black) presentation of ACV with visual target continuously oscillating after an initial 30 second baseline period. **c)** Tracking indices (TI) of eight wild type *D. erecta* males prior to (black), during (orange), and after (black) presentation of ACV with continuously oscillating visual target (black arrow) as in (b). **d)** and **e)** Left: Mean tracking index of wild type (d, n=8, same males shown in c) and anosmic *D. erecta* males mutant for the chemosensory co-receptors Orco and Ir8a (e, n=4), prior to (black), during (orange), and after (black) presentation of ACV with continuously oscillating visual target (black arrow). Thick lines represent the mean, shaded areas the standard error. Right: Probability of tracking. **f)** Left: Tracking index of *D. erecta* males expressing CsChrimson in Orco+ olfactory sensory neurons in the antenna and maxillary palps prior to (pre), during (LED) and after (post) optogenetic activation. Lines indicate mean ± standard error (shaded). Right: Probability of tracking. Those comparisons that were statistically significant are indicated as: * *p* ≤ 0.05

**Extended Data Figure 5.**
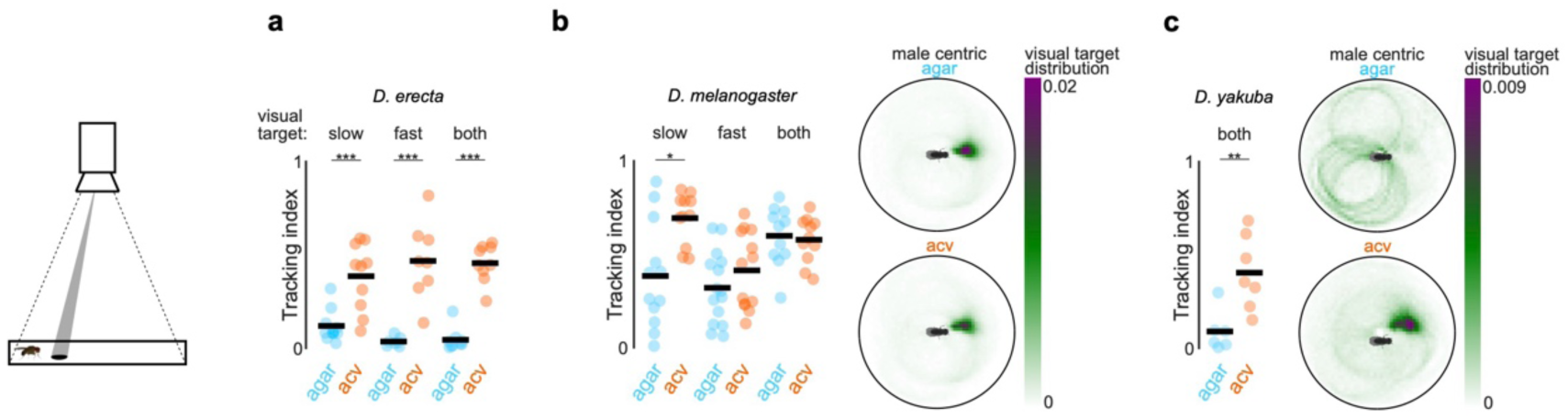
ACV plays a conserved role promoting courtship pursuit across species. **a)** Left: Tracking index of D. erecta towards either a slow (16 mm/sec) rotating dot, a fast (32 mm/sec) rotating dot, or both dots presented together in the presence of agar (blue) or ACV (orange). Data for the fast rotating dot is replotted from Fig. 3f. **b)** and **c)** Left: Tracking index of *D. melanogaster* (a) and *D. yakuba* (b) males towards either a slow (16 mm/sec) rotating dot, a fast (32 mm/sec) rotating dot, or both dots presented together in the presence of agar (blue) or ACV (orange). Right: Heatmap depicting the position of the fast visual target with respect to the egocentric orientation of the male (facing towards right) for assays in which both stimuli were present. Those comparisons that were statistically significant are indicated as: * *p* ≤ 0.05; \*\**p* < 0.001; *** *p* < 0.0001.

**Extended Data Figure 6.**
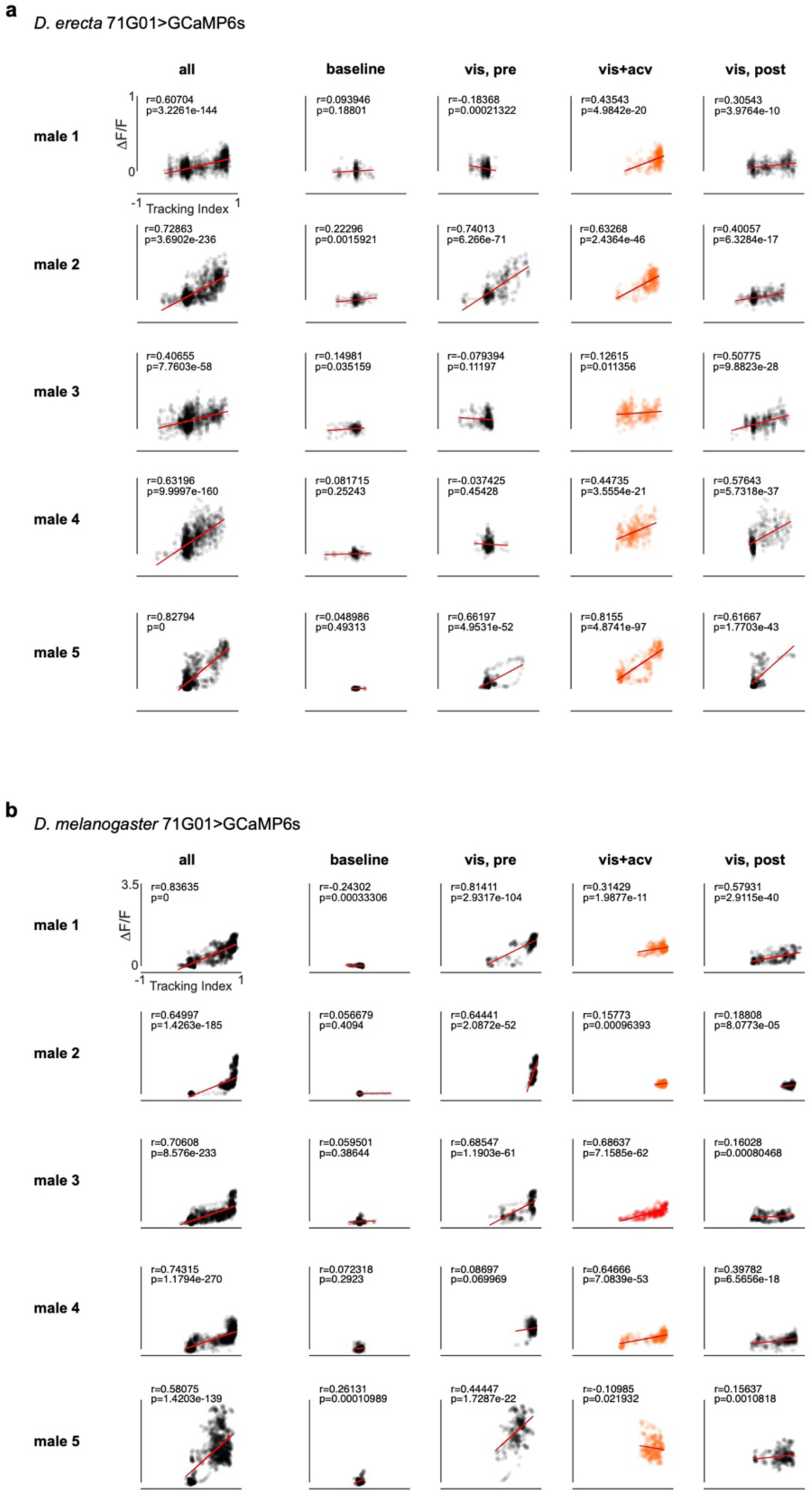
P1 neural activity is correlated with courtship tracking across species. **a)** and **b)** P1 neural activity (ΔF/F) of *D. erecta* (a) and *D. melanogaster* (b) males plotted against tracking index for five individuals (rows) for the entire experiment (all), during the 30 second baseline period (baseline), and with a continuously oscillating visual target prior to (vis, pre), during presentation of visual stimulus and ACV (vis + acv) and after (vis, post) presentation of ACV with only visual target. Line of best fit (red), Pearson correlation coefficient and p-value are indicated for each plot.

**Extended Data Figure 7.**
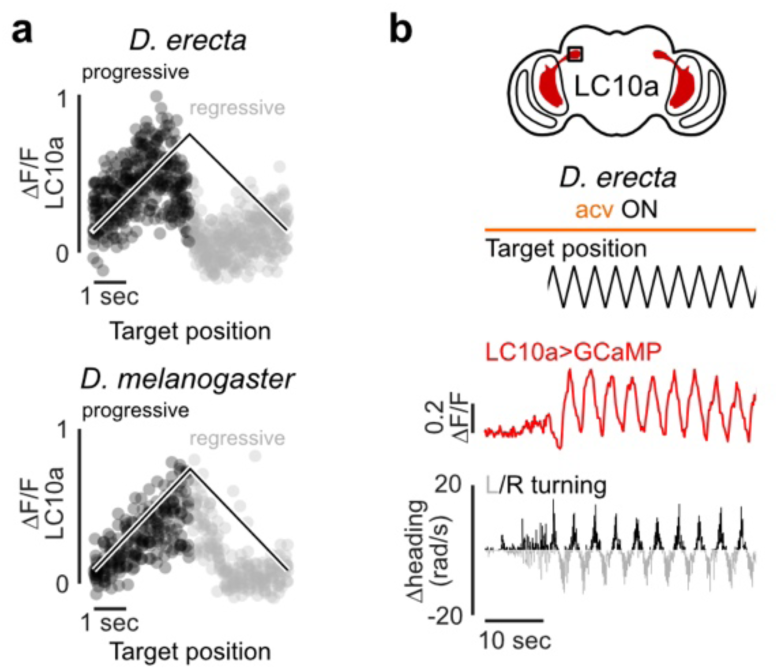
LC10a visual responses are similar in *D. erecta* and *D. melanogaster*. **a)** LC10a neural activity (ΔF/F) recorded in the right hemisphere of a representative *D. erecta* (above) and *D. melanogaster* (below) male tracking an oscillating visual target plotted against the position of the target (black line) in the male visual field colored by the target’s progressive (black) and regressive (grey) movement. ACV was provided throughout. **b)** Target position in the male visual field (top), corresponding LC10a neural activity (center), and heading direction (bottom) for a representative *D. erecta* male in the presence of ACV during an initial 10 second baseline period and then with visual target continuously oscillating.

**Extended Data Figure 8.**
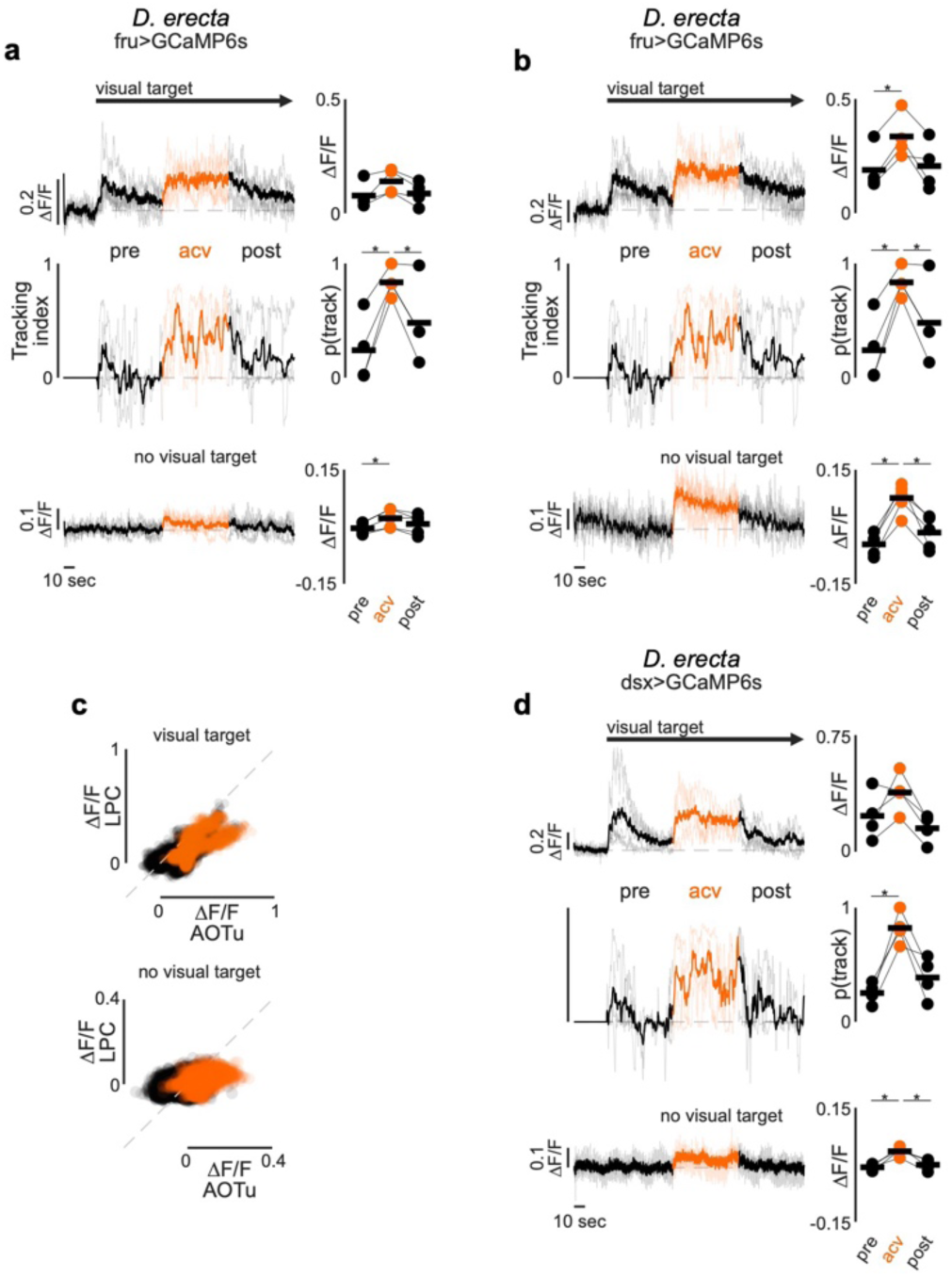
ACV activates distributed components of the courtship circuitry in *D. erecta*. **a)** Left: Neural activity (normalized ΔF/F) of aggregate Fruitless (Fru+) neurons in the lateral protocerebral complex (LPC) innervated by P1 neurons. Top: Neural activity. Center: tracking index of *D. erecta* males prior to (black), during (orange), and after (black) presentation of ACV with visual target continuously oscillating after an initial 30 second baseline period. Bottom: neural activity in the absence of a visual target. Thick lines represent the mean across individual males (thin lines). Arrow indicates appearance of visual target. Neural activity in LPC and AOTu was imaged simultaneously for each male. Right: Mean neural activity (top, bottom) and probability to track (center) before, during, and after presentation of ACV. **b)** Left: Neural activity (normalized ΔF/F) of aggregate Fruitless (Fru+) neurons in the anterior optic tubercle (AOTu) innervated by LC10a axons. Top: Neural activity. Center: tracking index of *D. erecta* males prior to (black), during (orange), and after (black) presentation of ACV with visual target continuously oscillating after an initial 30 second baseline period. Bottom: neural activity in the absence of a visual target. Thick lines represent the mean across individual males (thin lines). Arrow indicates appearance of visual target. Neural activity in LPC and AOTu was imaged simultaneously for each male. Right: Mean neural activity (top, bottom) and probability to track (center) before, during, and after presentation of ACV. **c)** Neural activity of Fru+ neurons in the lateral protocerebral complex (LPC) plotted against neural activity of Fru+ neurons in the anterior optic tubercle (AOTu) of *D. erecta* males shown in (a) and (b) with (top) and without (bottom) a continuously oscillating visual target. Dots represent individual frames color coded by the presence (orange) or absence (black) of ACV odor. Line of identity is indicated as grey dotted line. **d)** Left: Activity of Dsx+ neurons in the lateral protocerebral complex (LPC) innervated by P1 neurons (top) and tracking index (center) of *D. erecta* males prior to (black), during (orange), and after (black) presentation of ACV with visual target continuously oscillating after an initial 30 second baseline period and neural activity in the absence of a visual target (bottom). Neural activity is represented as normalized fluorescence (ΔF/F). Thick lines represent the mean across individual males (thin lines). Arrow indicates appearance of visual target. Right: Mean neural activity before, during, and after presentation of ACV (top, bottom) and probability to track (center). Those comparisons that were statistically significant are indicated as: * *p* ≤ 0.05

**Extended Data Figure 9.**
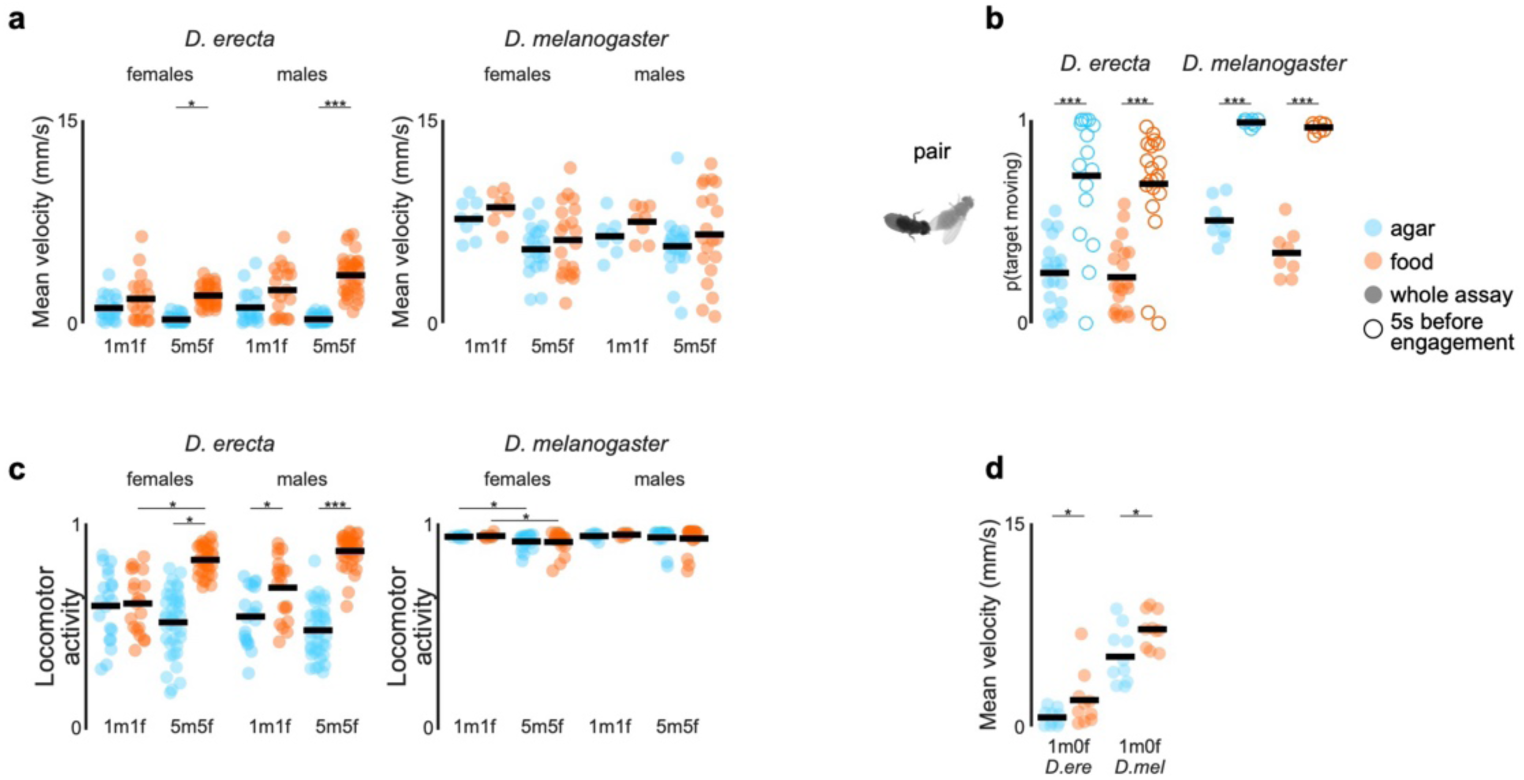
Locomotion in *D. erecta* depends on the chemosensory environment and scales with the group size. **a)** Mean velocity of individual female and male *D. erecta* (left) and *D. melanogaster* (right) in pairs (1m1f) or groups of 5 males and 5 females (5m5f) in small 38 mm arenas with a central 21.5 mm food patch formed from agar perfumed with ACV (orange) or an agar control (blue). Individual (dots) and overall means (black line) are indicated. **b)** Movement probability of a female target for *D. erecta* (left) and *D. melanogaster* (right) 1 male and 1 female pair assays in small 38 mm arenas with a central 21.5 mm food patch for the whole 40-minute assay (filled circle) and the five second period before the male engages in courtship (open circle) on agar (blue) or ACV (orange). Black line indicates mean. **c)** Fraction of time male (left) and female (right) *D. erecta* (D. ere) and *D. melanogaster* (D. mel) spent moving when paired (1m1f) or in groups (5m5f) over 40 minutes in a small 38 mm arena with a central 21.5 mm food patch on agar (blue) or ACV (orange). Black lines indicate mean. **d)** Mean velocity of individual *D. erecta* (left) and *D. melanogaster* (right) males in small 38 mm arenas with a central 21.5 mm food patch formed from agar perfumed with ACV (orange) or an agar control (blue). Individual (dots) and overall means (black line) are indicated. Those comparisons that were statistically significant are indicated as: * *p* ≤ 0.05; *** *p* < 0.0001.

**Extended Data Figure 10:**
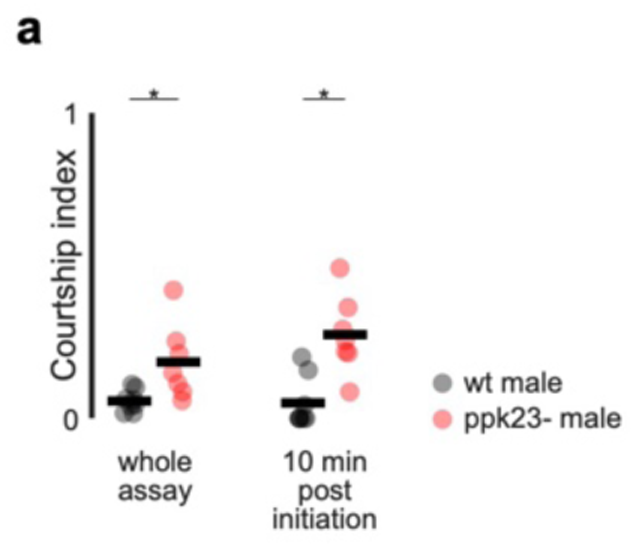
*D. erecta* remain averse to courting *D. melanogaster* females on ACV due to detection of heterospecific pheromones by ppk23+ sensory neurons. **a)** Courtship index of *D. erecta* males paired with a *D. melanogaster* female in small 38 mm arenas with a central 21.5 mm food patch formed from agar perfumed with ACV. Males were either wild type (black) or mutant for the DEG/ENaC sodium channel ppk23 (red). Courtship indices for the entire 40 minutes of the assay (left) or the 10 minutes following courtship initiation are shown. Black lines indicate mean. Enhanced courtship in mutants is consistent with evidence from other species that ppk23 is essential to detection of heterospecific pheromones. Those comparisons that were statistically significant are indicated as: * *p* ≤ 0.05

## Notes

### Competing Interest Statement

The authors have declared no competing interest.

## References

1. Bastock, M., & Manning, A. (1955). The courtship of *Drosophila melanogaster*. Behaviour, 85–111.

2. Carroll, C. R., & Janzen, D. H. (1973). Ecology of foraging by ants. Annual Review of Ecology and systematics, 231–257.

3. Hashikawa, K., Hashikawa, Y., Falkner, A., & Lin, D. (2016). The neural circuits of mating and fighting in male mice. Current opinion in neurobiology, 38, 27–37.

4. Anderson, D. J. (2016). Circuit modules linking internal states and social behaviour in flies and mice. Nature Reviews Neuroscience, 17(11), 692–704.

5. Severinghaus, L. L., Kurtak, B. H., & Eickwort, G. C. (1981). The reproductive behavior of Anthidium manicatum (Hymenoptera: Megachilidae) and the significance of size for territorial males. Behavioral Ecology and Sociobiology, 9(1), 51–58.

6. Yang, Z., Bengtsson, M., & Witzgall, P. (2004). Host plant volatiles synergize response to sex pheromone in codling moth, *Cydia pomonella*. Journal of chemical ecology, 30(3), 619–629.

7. Markow, T. A., & O’grady, P. (2008). Reproductive ecology of *Drosophila*. Functional Ecology, 747–759.

8. Hölldobler, B. (1976). The behavioral ecology of mating in harvester ants (Hymenoptera: Formicidae: Pogonomyrmex). Behavioral Ecology and Sociobiology, 405–423.

9. Allan, J. D., & Flecker, A. S. (1989). The mating biology of a mass-swarming mayfly. Animal Behaviour, 37, 361–371.

10. Rutowski, R. L. (1991). The evolution of male mate-locating behavior in butterflies. The American Naturalist, 138(5), 1121–1139.

11. Beani, L., Cervo, R., Lorenzi, C. M., & Turillazzi, S. (1992). Landmark-based mating systems in four *Polistes* species (Hymenoptera: Vespidae). Journal of the Kansas Entomological Society, 211–217.

12. Leboeuf, B. J. (1972). Sexual behavior in the northern elephant seal *Mirounga angustirostris*. Behaviour, 41(1-2), 1–26.

13. Slagsvold, T. (1986). Nest site settlement by the pied flycatcher: does the female choose her mate for the quality of his house or himself?. Ornis Scandinavica, 210–220.

14. Nelson, C. M. (1995). Male size, spawning pit size and female mate choice in a lekking cichlid fish. Animal Behaviour, 50(6), 1587–1599.

15. Paxton, R. J. (2005). Male mating behaviour and mating systems of bees: an overview. Apidologie, 36(2), 145–156.

16. Couzin, I. D. (2009). Collective cognition in animal groups. Trends in cognitive sciences, 13(1), 36–43.

17. Gibson, J. J. (2014). The ecological approach to visual perception: classic edition. Psychology press.

18. Maldonado-Chaparro, A. A., Montiglio, P. O., Forstmeier, W., Kempenaers, B., & Farine, D. R. (2018). Linking the fine-scale social environment to mating decisions: a future direction for the study of extra-pair paternity. Biological Reviews, 93(3), 1558–1577.

19. Sayin, S., Couzin-Fuchs, E., Petelski, I., Günzel, Y., Salahshour, M., Lee, C. Y., … & Couzin, I. D. (2025). The behavioral mechanisms governing collective motion in swarming locusts. Science, 387(6737), 995–1000.

20. Carson, H. L. (1971). The ecology of *Drosophila* breeding sites. The Harold L. Lyon Arboretum Lecture, University of Hawaii, 2, 1–27.

21. R’kha, S., Capy, P., & David, J. R. (1991). Host-plant specialization in the *Drosophila melanogaster* species complex: a physiological, behavioral, and genetical analysis. Proceedings of the National Academy of Sciences, 88(5), 1835–1839.

22. Mansourian, S., & Stensmyr, M. C. (2015). The chemical ecology of the fly. Current opinion in neurobiology, 34, 95–102.

23. Markow, T. A. (2019). Host use and host shifts in *Drosophila*. Current opinion in insect science, 31, 139–145.

24. Tsacas, L., Lachaise, D. (1974). Les Drosophilidae des savanes preforestleres de la region tropicale de Lamto (Cote-D’lvore). II. Le peuplement des fruits de *Pandanus candelabrum (Pandanacees*). In Annales de L’Universite D’Abidjan, Serie E (Vol. 7, pp. 153–192).

25. Rio, B., Couturier, G., Lemeunier, F., & Lachaise, D. (1983). Evolution d’une spécialisation saisonniére chez *Drosophila erecta* (Dipt., Drosophilidae). In Annales de la Société entomologique de France (NS) (Vol. 19, No. 2, pp. 235–248). Taylor & Francis.

26. Lachaise, D., Cariou, M. L., David, J. R., Lemeunier, F., Tsacas, L., & Ashburner, M. (1988). Historical biogeography of the *Drosophila melanogaster* species subgroup. In Evolutionary biology (pp. 159–225). Boston, MA: Springer US.

27. Dweck, H. K., Ebrahim, S. A., Kromann, S., Bown, D., Hillbur, Y., Sachse, S., … & Stensmyr, M. C. (2013). Olfactory preference for egg laying on citrus substrates in *Drosophila*. Current Biology, 23(24), 2472–2480.

28. Linz, J., Baschwitz, A., Strutz, A., Dweck, H. K., Sachse, S., Hansson, B. S., & Stensmyr, M. C. (2013). Host plant-driven sensory specialization in *Drosophila erecta*. Proceedings of the Royal Society B: Biological Sciences, 280(1760), 20130626.

29. Zhu, J., Park, K. C., & Baker, T. C. (2003). Identification of odors from overripe mango that attract vinegar flies, *Drosophila melanogaster*. Journal of chemical ecology, 29(4), 899–909.

30. Comeault, A. A., Serrato-Capuchina, A., Turissini, D. A., McLaughlin, P. J., David, J. R., & Matute, D. R. (2017). A nonrandom subset of olfactory genes is associated with host preference in the fruit fly Drosophila orena. Evolution Letters, 1(2), 73–85.

31. Bell, J. S., & Wilson, R. I. (2016). Behavior reveals selective summation and max pooling among olfactory processing channels. Neuron, 91(2), 425–438.

32. Kadakia, N., Demir, M., Michaelis, B. T., DeAngelis, B. D., Reidenbach, M. A., Clark, D. A., & Emonet, T. (2022). Odour motion sensing enhances navigation of complex plumes. Nature, 611(7937), 754–761.

33. Kohatsu, S., Koganezawa, M., & Yamamoto, D. (2011). Female contact activates male-specific interneurons that trigger stereotypic courtship behavior in *Drosophila*. Neuron, 69(3), 498–508.

34. Von Philipsborn, A. C., Liu, T., Jai, Y. Y., Masser, C., Bidaye, S. S., & Dickson, B. J. (2011). Neuronal control of *Drosophila* courtship song. Neuron, 69(3), 509–522.

35. Clowney, E. J., Iguchi, S., Bussell, J. J., Scheer, E., & Ruta, V. (2015). Multimodal chemosensory circuits controlling male courtship in *Drosophila*. Neuron, 87(5), 1036–1049.

36. Kallman, B. R., Kim, H., & Scott, K. (2015). Excitation and inhibition onto central courtship neurons biases Drosophila mate choice. Elife, 4, e11188.

37. Kohatsu, S., & Yamamoto, D. (2015). Visually induced initiation of *Drosophila* innate courtship-like following pursuit is mediated by central excitatory state. Nature communications, 6(1), 6457.

38. Jallon, J. M., & David, J. R. (1987). Variations in cuticular hydrocarbons among the eight species of the *Drosophila melanogaster* subgroup. Evolution, 41(2), 294–302.

39. Khallaf, M. A., Cui, R., Weißflog, J., Erdogmus, M., Svatoš, A., Dweck, H. K., … & Knaden, M. (2021). Large-scale characterization of sex pheromone communication systems in *Drosophila*. Nature communications, 12(1), 4165.

40. Coleman, R. T., Morantte, I., Koreman, G. T., Cheng, M. L., Ding, Y., & Ruta, V. (2024). A modular circuit coordinates the diversification of courtship strategies. Nature, 635(8037), 142–150.

41. Hoopfer, E. D., Jung, Y., Inagaki, H. K., Rubin, G. M., & Anderson, D. J. (2015). P1 interneurons promote a persistent internal state that enhances inter-male aggression in *Drosophila*. Elife, 4, e11346.

42. Hindmarsh Sten, T., Li, R., Otopalik, A., & Ruta, V. (2021). Sexual arousal gates visual processing during *Drosophila* courtship. Nature, 595(7868), 549–553.

43. Schretter, C. E., Hindmarsh Sten, T., Klapoetke, N., Shao, M., Nern, A., Dreher, M., … & Rubin, G. M. (2025). Social state alters vision using three circuit mechanisms in *Drosophila*. Nature, 637(8046), 646–653.

44. Ribeiro, I. M., Drews, M., Bahl, A., Machacek, C., Borst, A., & Dickson, B. J. (2018). Visual projection neurons mediating directed courtship in Drosophila. Cell, 174(3), 607–621.

45. Barrows, W. M. (1907). The Reactions of the Pomace Fly: *Drosophila ampelophila* Loew, to Odorous Substances.

46. Da Cunha, A. B., Dobzhansky, T., & Sokoloff, A. (1951). On food preferences of sympatric species of *Drosophila*. Evolution, 97–101.

47. Grosjean, Y., Rytz, R., Farine, J. P., Abuin, L., Cortot, J., Jefferis, G. S., & Benton, R. (2011). An olfactory receptor for food-derived odours promotes male courtship in *Drosophila*. Nature, 478(7368), 236–240.

48. Cheng, K. Y., Colbath, R. A., & Frye, M. A. (2019). Olfactory and neuromodulatory signals reverse visual object avoidance to approach in *Drosophila*. Current Biology, 29(12), 2058–2065.

49. Cheng, K. Y., & Frye, M. A. (2021). Odour boosts visual object approach in flies. Biology Letters, 17(3), 20200770.

50. Seeholzer, L. F., Seppo, M., Stern, D. L., & Ruta, V. (2018). Evolution of a central neural circuit underlies *Drosophila* mate preferences. Nature, 559(7715), 564–569.

51. Ye, D., Walsh, J. T., Junker, I. P., & Ding, Y. (2024). Changes in the cellular makeup of motor patterning circuits drive courtship song evolution in *Drosophila*. Current Biology, 34(11), 2319–2329.

52. Partridge, L., Hoffmann, A., & Jones, J. S. (1987). Male size and mating success in *Drosophila melanogaster* and *D. pseudoobscura* under field conditions. Animal Behaviour, 35(2), 468–476.

53. Garbaczewska, M., Billeter, J. C., & Levine, J. D. (2013). *Drosophila melanogaster* males increase the number of sperm in their ejaculate when perceiving rival males. Journal of insect physiology, 59(3), 306–310.

54. Dukas, R. (2020). Natural history of social and sexual behavior in fruit flies. Scientific reports, 10(1), 21932.

55. Hindmarsh Sten, T. A., Li, R., Hollunder, F., Eleazer, S., & Ruta, V. (2025). Male-male interactions shape mate selection in *Drosophila*. Cell, 188(6), 1486–1503.

56. Katz, Y., Tunstrøm, K., Ioannou, C. C., Huepe, C., & Couzin, I. D. (2011). Inferring the structure and dynamics of interactions in schooling fish. Proceedings of the National Academy of Sciences, 108(46), 18720–18725.

57. Ballerini, M., Cabibbo, N., Candelier, R., Cavagna, A., Cisbani, E., Giardina, I., … & Zdravkovic, V. (2008). Empirical investigation of starling flocks: a benchmark study in collective animal behaviour. Animal behaviour, 76(1), 201–215.

58. Cavagna, A., Cimarelli, A., Giardina, I., Parisi, G., Santagati, R., Stefanini, F., & Viale, M. (2010). Scale-free correlations in starling flocks. Proceedings of the National Academy of Sciences, 107(26), 11865–11870.

59. Larsch, J., & Baier, H. (2018). Biological motion as an innate perceptual mechanism driving social affiliation. Current Biology, 28(22), 3523–3532.

60. Sridhar, V. H., Li, L., Gorbonos, D., Nagy, M., Schell, B. R., Sorochkin, T., … & Couzin, I. D. (2021). The geometry of decision-making in individuals and collectives. Proceedings of the National Academy of Sciences, 118(50), e2102157118.

61. Li, L., Nagy, M., Amichay, G., Wu, R., Wang, W., Deussen, O., … & Couzin, I. D. (2025). Reverse engineering the control law for schooling in zebrafish using virtual reality. Science Robotics, 10(101), eadq6784.

62. Stockinger, P., Kvitsiani, D., Rotkopf, S., Tirián, L., & Dickson, B. J. (2005). Neural circuitry that governs *Drosophila* male courtship behavior. Cell, 121(5), 795–807.

63. Simon, J. C., & Dickinson, M. H. (2010). A new chamber for studying the behavior of *Drosophila*. Plos one, 5(1), e8793.

64. Eyjolfsdottir, E., Branson, S., Burgos-Artizzu, X. P., Hoopfer, E. D., Schor, J., Anderson, D. J., & Perona, P. (2014, September). Detecting social actions of fruit flies. In European Conference on Computer Vision (pp. 772–787). Cham: Springer International Publishing.

65. Chiara, V., & Kim, S. Y. (2023). AnimalTA: A highly flexible and easy-to-use program for tracking and analysing animal movement in different environments. Methods in Ecology and Evolution, 14(7), 1699–1707.

66. Friard, O., & Gamba, M. (2016). BORIS: a free, versatile open-source event-logging software for video/audio coding and live observations. Methods in ecology and evolution, 7(11), 1325–1330.

67. Ryba, A., Brand, P., Coleman, R. T., Greenfeld, Y., Tsitohay, Y. N., Hollunder, F., … & Ruta, V. (2025). Strain variation identifies a neural substrate for behavioral evolution in Drosophila. bioRxiv, 2025-08.

68. Maimon, G., Straw, A. D., & Dickinson, M. H. (2010). Active flight increases the gain of visual motion processing in Drosophila. Nature neuroscience, 13(3), 393–399.

